# RNase L Regulates Antiviral Responsiveness through Cleavage of XBP1 mRNA

**DOI:** 10.64898/2026.03.21.713401

**Authors:** Yoshika Takenaka, Yasutoshi Akiyama, Tsubasa Inaba, Daiki Shinozuka, Katsuki Aoyama, Ransu Ogasawara, Nana Kunii, Takaaki Abe, Eiji Morita, Yoshihisa Tomioka, Pavel Ivanov

## Abstract

During viral infection, viral replication perturbs endoplasmic reticulum (ER) homeostasis and triggers the unfolded protein response (UPR). XBP1s, a transcription factor generated by one branch of the UPR, is known to potentiate both innate and adaptive immunity, but its role in antiviral responses remains incompletely understood beyond its ability to augment type I interferon (IFN) mRNA induction. Here, we show that XBP1s positively regulates the RIG-I-like receptors (RLRs), ribonuclease L (RNase L), and protein kinase R (PKR) pathways, indicating that it enhances all three major antiviral response pathways. We further show that RNase L activation rapidly decreases XBP1 mRNA levels in an RNase activity-dependent manner, leading to a prompt reduction in XBP1s expression. Consistent with this, RNase L deletion significantly increased both thapsigargin-mediated XBP1s induction and XBP1s expression following Japan encephalitis virus infection. Poly(I:C)-induced IFNB mRNA expression was significantly enhanced in RNase L-knockout cells. This enhancement was completely abolished by RNase L reconstitution. XBP1 knockdown also significantly attenuated IFNB mRNA expression in RNase L-knockout cells. These findings suggest a negative-feedback loop in which RNase L suppresses XBP1s, thereby fine-tuning antiviral responsiveness during viral infection.

**GRAPHICAL ABSTRACT:** 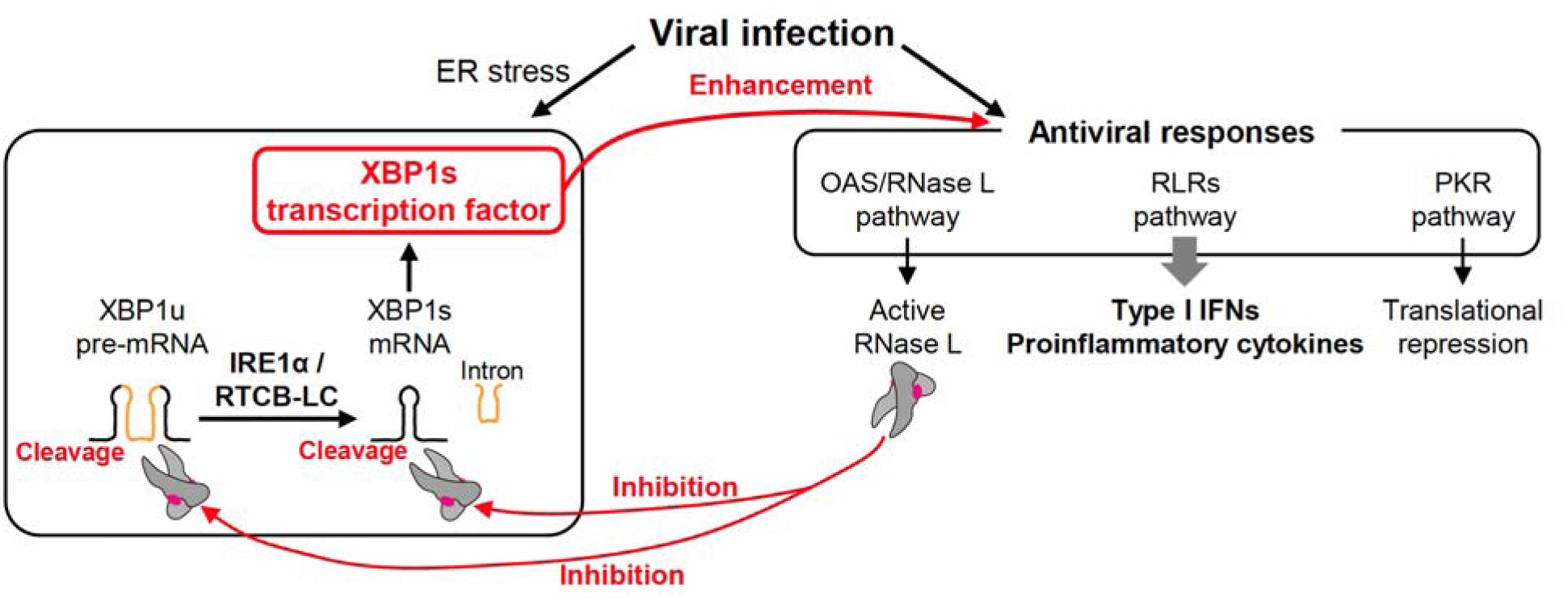

## INTRODUCTION

The innate immune system provides the first line of defense against invading pathogens and mediates the rapid initial response to infection. Through pattern recognition receptors (PRRs), it detects conserved microbial features and generates signals that not only initiate immediate antimicrobial defenses but also shape subsequent adaptive immune responses. (1,2). During viral infection, viral replication generates intracellular double-stranded RNA (dsRNA), which is recognized by cellular dsRNA sensors and initiates antiviral responses. Among the best-characterized dsRNA-triggered pathways are the retinoic acid-inducible gene I (RIG-I)-like receptors (RLRs), protein kinase R (PKR), and ribonuclease L (RNase L) pathways (3,4). The RLR pathway is initiated when RIG-I and melanoma differentiation-associated protein 5 (MDA5) recognize viral dsRNA and signal through mitochondrial antiviral-signaling protein (MAVS), thereby inducing the production of type I interferons (IFNs) and proinflammatory cytokines. In contrast, the PKR pathway is activated by dsRNA-bound PKR, which phosphorylates the α subunit of eukaryotic initiation factor 2 (eIF2α) and inhibits translation initiation as a branch of the integrated stress response (ISR) (5,6). In the RNase L pathway, double-stranded RNA (dsRNA) is sensed by 2’-5’-oligoadenylate synthetases (OASs), which synthesize 2’-5’-linked oligoadenylates (2-5A) that activate RNase L (7). Activated RNase L then cleaves viral and host RNAs, thereby contributing to the antiviral response (8). Recent studies have revealed that RNase L contributes to antiviral defense through multiple mechanisms beyond direct RNA degradation. In addition to promoting global translational shutoff through widespread host mRNA decay (9,10), RNase L can trigger a ZAKα-dependent ribotoxic stress response leading to proinflammatory signaling and apoptosis (11,12), and can also remodel nuclear RNA biogenesis by altering mRNA processing and promoting nuclear retention of antiviral transcripts such as IFN-β (13). Moreover, RNase L-generated RNA fragments have been reported to amplify innate immune signaling through RLR-dependent pathways (14). In addition, RNase L has been reported to cleave a broad range of host RNAs, including rRNAs and tRNAs (15). In the case of tRNAs, RNase L-mediated in the anticodon loop generates a class of small RNA known as tRNA-derived RNAs (tDRs) (16,17). We recently reported that RNase L cleaves multiple tRNA species to produce tDRs, and that the resulting 5’-tDR^Ala^ exerts a translation inhibitory effect, suggesting that RNase L may also contribute to the stress response during viral infection through tDR production (18).

It is well established that viral infection imposes stress on the endoplasmic reticulum (ER), thereby activating the unfolded protein response (UPR) (19,20). This is thought to result not only from the increased burden associated with viral protein synthesis, folding, and processing, but also from broader perturbations of ER homeostasis during infection. The UPR comprises three signaling branches: the inositol-requiring enzyme 1α (IRE1α)-X-box binding protein 1 (XBP1) branch, the activating transcription factor 6 (ATF6) branch, and the protein kinase RNA-like endoplasmic reticulum kinase (PERK) branch (21). In the IRE1α-XBP1 branch, the ER-resident transmembrane protein IRE1α cleaves unspliced XBP1 (XBP1u) pre-mRNA, after which the resulting exons are ligated by the RNA ligase RTCB complex (RTCB-LC) to generate spliced XBP1 (XBP1s) mRNA (22–24). Translation of XBP1s mRNA produces the transcription factor XBP1s, which enhances the expression of UPR-related genes (25). XBP1s is well known not only to promote the expression of genes involved in ER protein folding, but also to broadly potentiate immune responses in both the innate and adaptive immune systems (26). In innate immunity, XBP1s has been reported to promote proinflammatory cytokine production in macrophages (27) and to support the development and survival of dendritic cells (28). However, although viral infection is widely known to induce ER stress and activate the UPR, the role of XBP1s in antiviral responses remains largely unclear, except for its enhancing effect on the RLR pathway through promotion of IFNB mRNA induction (29,30).

In addition to catalyzing XBP1 mRNA splicing, the RTCB-LC also mediates precursor tRNA (pre-tRNA) splicing (31). Furthermore, the RTCB-LC also contributes to the regulation of tDR production. Angiogenin (ANG) is a stress-responsive RNase that becomes activated under stress and cleaves mature tRNAs at the anticodon loop, thereby generating tDRs termed tiRNAs (tRNA-derived stress-induced RNAs) (17,32). Notably, 5’-tiRNA^Ala^ and 5’-tiRNA^Cys^ have been shown to inhibit translation (33), supporting their roles as stress-responsive effectors (34). We recently reported that the RTCB-LC controls tiRNA levels by repairing ANG-cleaved tiRNAs (35). We further showed that, because the RTCB-LC is inhibited under oxidative stress (36), this repair process is suppressed under such conditions, thereby enhancing tiRNA production (35). These findings suggest that the RTCB-LC functions, particularly during oxidative stress, to promote the accumulation of tiRNAs, which act as stress-responsive molecules. Enzymatically, the RTCB-LC ligates RNA ends containing a 2’,3’-cyclic phosphate and a 5’-hydroxyl group (37,38). Since RNase L generates RNA cleavage products with these termini (15), RNase L-induced tDRs, including 5’-tDR^Ala^ that shows a translation inhibitory effect (18), are also predicted to be potential substrates of the RTCB-LC and may therefore be regulated in a manner similar to ANG-generated tiRNAs. However, it remains unknown whether the RTCB-LC indeed regulates the levels of RNase L-induced tDRs.

In this study, we first show that inhibition of the RTCB-LC increases the production of RNase L-induced tDRs, indicating that the RTCB-LC negatively regulates RNase L-dependent tDR generation, as is the case for ANG-induced tiRNAs. In contrast, prolonged suppression of the RTCB-LC by knockdown reduced the production of RNase L-induced tDRs. From these seemingly contradictory results, we revealed that XBP1s positively regulates the expression of RNase L and OAS family members, and that RTCB also contributes to the responsiveness of the RNase L pathway through its control of XBP1s production. Further investigation revealed that XBP1s positively regulates the responsiveness of all three major antiviral response pathways. We further demonstrate that RNase L, once activated, rapidly suppresses XBP1s production through cleavage of XBP1 mRNA, thereby limiting the XBP1s-dependent enhancement of type I IFN induction. This inhibitory pathway suggests a potential negative feedback mechanism by which RNase L restrains the immunoenhancing activity of XBP1s. Together, our findings reveal a previously unrecognized mechanism by which RNase L modulates antiviral responses.

## MATERIAL AND METHODS

### Cell Culture and Treatment

Two human cell lines, U2OS and A549, were used in this study. RNASEL KO (ΔRNASEL) A549 cells were generated as previously described (18). Japanese Encephalitis Virus (JEV) stocks were prepared using 293T and Vero cells. Cells were maintained at 37 °C in 5% CO_2_ incubator in Dulbecco’s modified Eagle’s medium (DMEM) (Nacalai Tesque) supplemented with 10% fetal bovine serum (Corning) and 1% of penicillin-streptomycin mixed solution (Nacalai Tesque).

For oxidative inhibition of RTCB, hydrogen peroxide (Nacalai Tesque) was used at 1 mM for 2 hours. To induce ER stress, cells were treated with tunicamycin (FUJIFILM Wako Pure Chemical Corporation) at a final concentration of 2 μg/mL for 16 hours, or with thapsigargin (FUJIFILM Wako Pure Chemical Corporation) at 25 nM for the indicated times. To induce antiviral responses, cells were transfected with Poly(I:C) (HMW) (InvivoGen) at a final concentration of 1 or 2 µg/mL using Lipofectamine 3000 (ThermoFisher) according to the manufacturer’s protocol. When Poly(I:C) (InvivoGen) was used alone, cells were transfected with Poly(I:C) at a final concentration of 2 µg/mL for 6 hours unless otherwise indicated. In the combined Poly(I:C) transfection and thapsigargin-induced stress model, Poly(I:C) was transfected at 1 µg/mL, and thapsigargin was administered at 25 nM. Cells were collected using RIPA buffer for protein purification or Sepasol-RNA I Super G (Nacalai Tesque) for RNA purification.

### Western blotting

Western blotting was performed as we previously reported (18,35). Antibodies used in this study are shown in Supplemental Table 1.

### Plasmids

DNA oligonucleotides for shRNA targeting RTCB or XBP1 were designed using siDirect (39), synthesized by Integrated DNA Technologies (IDT), and subcloned into pLKO.1 - TRC cloning vector (Addgene) according to the manufacturer’s instructions. The sequences of the oligonucleotides are shown in Supplemental Table 2. pLKO.1 - TRC cloning vector was a gift from David Root (Addgene plasmid # 10878).

FLAG-tagged RNASEL was amplified by polymerase chain reaction (PCR) using complementary DNA (cDNA) synthesized from total RNA isolated from U2OS cells. The PCR amplicon was then subcloned into the pcDNA3.1 (+) vector (ThermoFisher) using BamHI and EcoRI restriction sites. To disrupt the PAM sequence and prevent Cas9-mediated cleavage, a silent mutation was introduced into the coding sequence (CDS) by site-directed mutagenesis using KOD -Plus- Mutagenesis Kit (TOYOBO), changing the 20th codon from AGG to AGA. A catalytically inactive R667A mutant was generated by substitution of CGG to GCC. To generate a lentiviral construct for RNase L expression, the RNASEL CDS (WT/R667A mutant) was subcloned into the FUGW vector (Addgene). The sequences of the PCR primers are shown in Supplemental Table 3. FUGW was a gift from David Baltimore (Addgene plasmid # 14883).

For XBP1s overexpression, the CDS of human XBP1s was amplified by PCR, then subcloned into between BamH I and Xho I site of the pcDNA3.1 (+) vector (ThermoFisher). To facilitate PCR amplification of XBP1s mRNA, cDNA was prepared using total RNA extracted from tunicamycin-treated U2OS cells. The sequences of the PCR primers are shown in Supplemental Table 3.

### Generation of lentiviruses

Lentiviral particles were generated as we previously reported (18,35).

### Knockdown of RTCB/XBP1

Knockdown was performed as we previously reported (18,35). Knockdown efficiencies of RTCB or XBP1 were evaluated by Western blotting. Western blotting against anti-ACTB antibody (Proteintech) was used as loading control. Ponceau S staining (Nacalai Tesque) was performed to verify equal loading.

### Generation of ΔRNASEL_FLAG-RNASEL (WT/R667A) Cells

The lentiviral particles were transduced into ΔRNASEL cells in the presence of 8 μg/mL polybrene (Sigma-Aldrich). Cells were cloned by limiting dilution, then expression of FLAG-RNase L in individual clones was confirmed by Western blotting.

### Overexpression of XBP1s

Two microgram of the pcDNA3.1 (+) (Mock) or pcDNA3.1_XBP1s was transfected into U2OS cells (5 x 10^5^ cells) in a 6-well dish using Lipofectamine 3000 (ThermoFisher) according to the manufacturer’s protocol. Twenty-four hours after transfection, cells were collected with Sepasol-RNA I Super G (Nacalai Tesque) for RNA purification.

### Northern Blotting

Northen Blotting was performed as we previously reported (18,35,40). The sequences of the probes used in this study are shown in Supplemental Table 4.

### Quantitative PCR (qPCR)

Complementary DNA (cDNA) was synthesized from 1 µg of total RNA using High-Capacity cDNA Reverse Transcription Kit (Applied Biosystems) according to the manufacturer’s instructions. qPCR was performed using THUNDERBIRD SYBR qPCR Mix (TOYOBO) according to the manufacturer’s instructions using CFX Connect Real-Time System (Bio-Rad Laboratories). Unless otherwise indicated, relative mRNA levels were calculated from Cq values. When indicated, relative mRNA expression levels were calculated using the ΔΔCt method, with GAPDH mRNA was used as an internal control. The sequences of the probes used in this study are shown in Supplemental Table 5.

### JEV infection Assay

A recombinant Japanese encephalitis virus (JEV) encoding an NS1 protein fused to a HiBiT tag was generated using circular polymerase extension reaction (CPER) as previously described (41). To construct the viral genome, six cDNA fragments spanning the complete genome of the JEV strain AT31 (GenBank accession number AB196926) were first cloned into pUC19 vectors. Each fragment was then amplified by PCR and assembled via CPER. The resulting CPER products were transfected into 293T cells using polyethylenimine Max (PEI MAX) (Polyscience, Warrington, PA, USA). Culture supernatants were collected at 6 days post-transfection, and viral titers were determined by focus-forming assay using Vero cells. For subsequent experiments, cells were infected with the recombinant virus at a multiplicity of infection (MOI) of 0.3. Viral replication was assessed by measuring the amount of HiBiT-fused NS1 protein released into the culture supernatant, quantified using a HiBiT-dependent NanoLuc luciferase assay.

### Statistical Analyses

All data are presented as mean ± SD. Comparisons between two experimental groups were performed using Student’s t-test. For multiple group comparisons, one-way ANOVA followed by Tukey–Kramer post hoc test was used. To analyze the interaction between two factors, two-way ANOVA was conducted. All statistical analyses were performed using JMP Student Edition (JMP Statistical Discovery) or Kyplot 6.0 (KyensLab).

## RESULTS

### RTCB complex regulates RNase L-induced tDR production

We recently reported that RNase L cleaves various tRNA species into tDRs, including 5’-tDR^Ala^ which shows a translational inhibitory effect (18), suggesting that RNase L-mediated tDR production may function as a stress response during viral infection. Like ANG, RNase L is a metal-independent RNase and therefore generates 5’-tDRs bearing a 2’,3’-cyclic phosphate and 3’-tDRs with a 5’-hydroxyl group (42), which together constitute substrates for RTCB-LC. We therefore sought to determine whether RNase L-mediated tDR production, as is the case of ANG-mediated tiRNA production, is regulated by RTCB-LC. To address this question, we examined whether inhibition of RTCB-LC increased the amount of RNase L-induced tDRs (Figure 1A).

**Figure 1:**
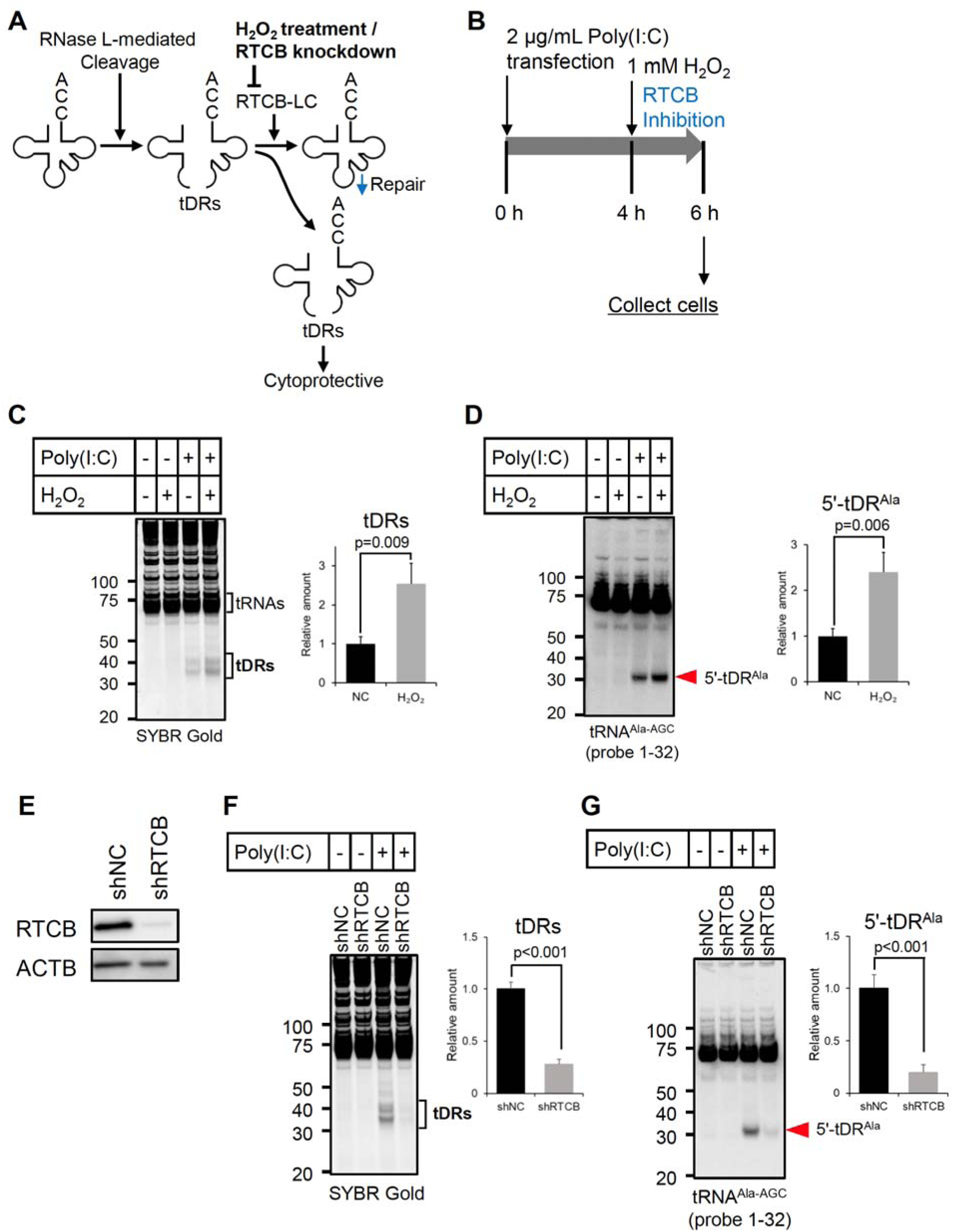
RTCB complex regulates RNase L-induced tDR production. (A) Model proposing that inhibition of RTCB function by H_2_O_2_ or RTCB knockdown reduces RTCB complex-dependent tDR repair, resulting in elevated levels of RNase L-induced tDRs. (B-D) H_2_O_2_ treatment boosts RNase L-induced tDR production in U2OS cells. (B) A schematic diagram of the experimental design. (C) SYBR Gold staining and (D) Northern blotting for tRNA^Ala-AGC^. Relative amounts of tDRs and 5’-tDR^Ala^ calculated by densitometry are also shown (n=3 each). (E-G) RTCB knockdown significantly decreases the amount of RNase L-induced tDRs in U2OS cells. (E) RTCB protein levels 7 days after shRNA-mediated RTCB knockdown. Western blotting for RTCB and ACTB are shown. (F) SYBR Gold staining and (G) Northern blotting for tRNA^Ala-AGC^. Relative amounts of tDRs and 5’-tDR^Ala^ calculated by densitometry are also shown (n=4 each).

We first examined whether a 2-hour H_2_O_2_ treatment, which suppresses the enzymatic activity of RTCB-LC (36) while not itself inducing ANG-mediated tiRNA production (35), affects RNase L-induced tDR levels in U2OS cells (Figure 1B). As shown in Figure 1C and 1D, H_2_O_2_ itself did not induce tDR production, suggesting that a 2-h H_2_O_2_ treatment did not activate stress responsive RNases including ANG. Under the condition, the amount of RNase L-induced tDRs including 5’-tDR^Ala^ was significantly increased by H_2_O_2_ treatment (Figure 1C, D). In A549 cells, RNase L-induced tDR levels were also significantly increased by H_2_O_2_ treatment (Supplemental Fig. S1). These data suggest that as with ANG-generated tiRNAs, RTCB-LC negatively regulates RNase L-induced tDR production, likely the same mechanism in which RTCB-LC mediates the repair of RNase L-cleaved tDRs by ligating the 5’- and 3’-tDRs prior to their dissociation (35). We next examined whether knockdown of RTCB, as with H_2_O_2_ treatment, increases the levels of RNase L-generated tDRs (Figure 1E-G). Unexpectedly, in contrast to H_2_O_2_ treatment, the levels of RNase L-induced tDRs, including 5’-tDR^Ala^, were significantly decreased by RTCB knockdown (Figure 1F, G).

### XBP1s acts as a positive regulator of the RNase L pathway

What underlies the contrasting outcomes of H_2_O_2_ treatment and RTCB knockdown? Given that RTCB-LC is responsible for the production of XBP1s transcription factor (22–24), we considered the possibility that XBP1s regulates the expression of genes involved in the RNase L pathway, including RNase L itself (Figure 2A). To investigate the impact of XBP1s transcription factor on RNase L pathway, we first examined whether RTCB knockdown decreased XBP1s production. Since XBP1s was not detectable under basal conditions by Western blotting, we evaluated XBP1s levels under ER stress conditions induced by tunicamycin (Figure 2B). RTCB/XBP1 knockdown significantly inhibited tunicamycin-induced XBP1s production both in U2OS and A549 cells (Figure 2B, Supplemental Fig. S2A). RNase L levels were significantly decreased by either RTCB or XBP1 knockdown (Figure 2C, Supplemental Fig. S2B), whereas overexpression of XBP1s significantly increased RNASEL mRNA levels (Figure 2D), which suggests that XBP1s positively regulates RNase L expression. We also investigated the expression levels of 2’-5’-oligoadenylate synthetases (OASs), which are activated by binding to dsRNAs and subsequently activate RNase L (7). Western blotting showed that OAS3, an OAS family member primarily responsible for RNase L activation (43), was significantly reduced by RTCB/XBP1 knockdown (Figure 2E). While OAS2 expression could not be reliably quantified due to its low abundance, mRNA levels of OAS1 and OAS3 were significantly decreased by RTCB/XBP1 knockdown (Figure 2F). Conversely, XBP1s overexpression significantly increased mRNA levels of OAS1 and OAS3 (Figure 2G). These results indicate that XBP1s positively regulates not only RNase L but also the expression of OASs, which function as activators of RNase L, suggesting that XBP1s plays a central role in controlling the responsiveness of RNase L pathway. Similar to RTCB knockdown, U2OS cells showed a significant decrease in RNase L-induced tDR levels, including 5’-tDR^Ala^ (Figure 2H, I) upon XBP1 knockdown, while the decrease in A549 cells did not reach statistical significance (Supplemental Fig. S2C, D). Given that RNASEL/OAS3 knockdown decreases Poly(I:C)-induced tDR production likely due to a reduction in the overall levels of active RNase L (18), these results indicate that the reduction in Poly(I:C)-induced tDRs observed under RTCB/XBP1 knockdown is likely due to decreased expression of RNase L and OASs, thereby reducing RNase L-mediated tRNA cleavage. Taken together, these data suggest that XBP1s determines the magnitude of the RNase L pathway under viral infection.

**Figure 2:**
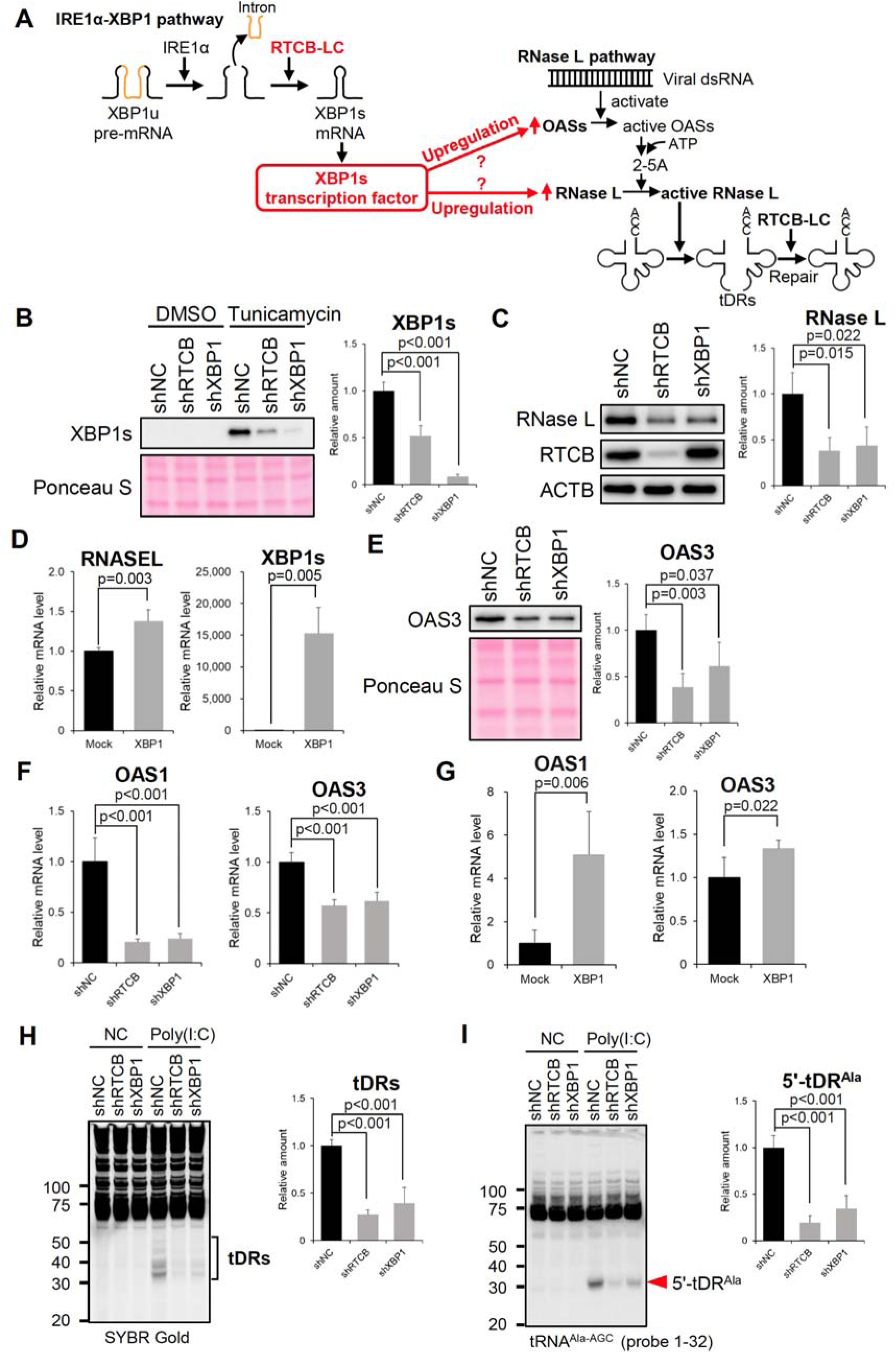
XBP1s acts as a positive regulator of the RNase L pathway. (A) A hypothetical model for the regulation of the RNase L pathway by XBP1s transcription factor. (B) RTCB/XBP1 knockdown significantly decreases tunicamycin-induced XBP1s production. Relative amounts of tunicamycin-induced XBP1s calculated by densitometry are also shown (n=4). (C, D) XBP1s positively regulates RNase L expression in U2OS cells. (C) RTCB/XBP1 knockdown significantly decreases RNase L expression. Western blotting for RNase L, RTCB and ACTB are shown. Relative amounts of RNase L calculated by densitometry are also shown (n=3). (D) XBP1s overexpression significantly increases RNase L expression. qPCR for RNASEL and XBP1s. Relative expression levels to GAPDH mRNA are shown. Means and standard deviation were obtained from four independent experiments. (E-G) XBP1s also upregulates OAS expression in U2OS cells. (E) RTCB/XBP1 knockdown significantly decreases OAS3 expression. Western blotting for OAS3. The amounts of OAS3 calculated by densitometry are also shown (n=4). (F) RTCB knockdown significantly decreases OAS1 and OAS3 mRNA expression. qPCR for OAS1 and OAS3. Relative expression levels to GAPDH mRNA are shown. Means and standard deviation were obtained from four independent experiments. (G) XBP1s overexpression significantly increases OAS1 and OAS3 expression. qPCR for OAS1 and OAS3. Relative expression levels to GAPDH mRNA are shown. Means and standard deviation were obtained from four independent experiments. (H, I) RTCB/XBP1 knockdown decreases RNase L-induced tDRs in U2OS cells. (H) SYBR Gold staining and (I) Northern blotting for tRNA^Ala-AGC^. Relative amount of tDRs and 5’-tDR^Ala^ calculated by densitometry are also shown (n=4 each).

### XBP1s positively regulates all three antiviral response pathways

Following the finding that transcription factor XBP1s positively regulates the expression of RNase L pathway genes, we next asked whether XBP1s also controls the expression of genes in the RLR and PKR pathways (Figure 3A and Supplemental Fig. S3A). With respect to the RLR pathway, expression of the dsRNA receptor MDA5 was significantly reduced by RTCB/XBP1 knockdown, whereas expression of the downstream adaptor molecule MAVS remained unchanged. In the PKR pathway, expression levels of both PKR and PACT, a regulator of PKR (3,44), were significantly decreased following RTCB/XBP1 knockdown. Consistent results were obtained in both U2OS and A549 cells (Figure 3A and Supplemental Fig. S3A).

**Figure 3:**
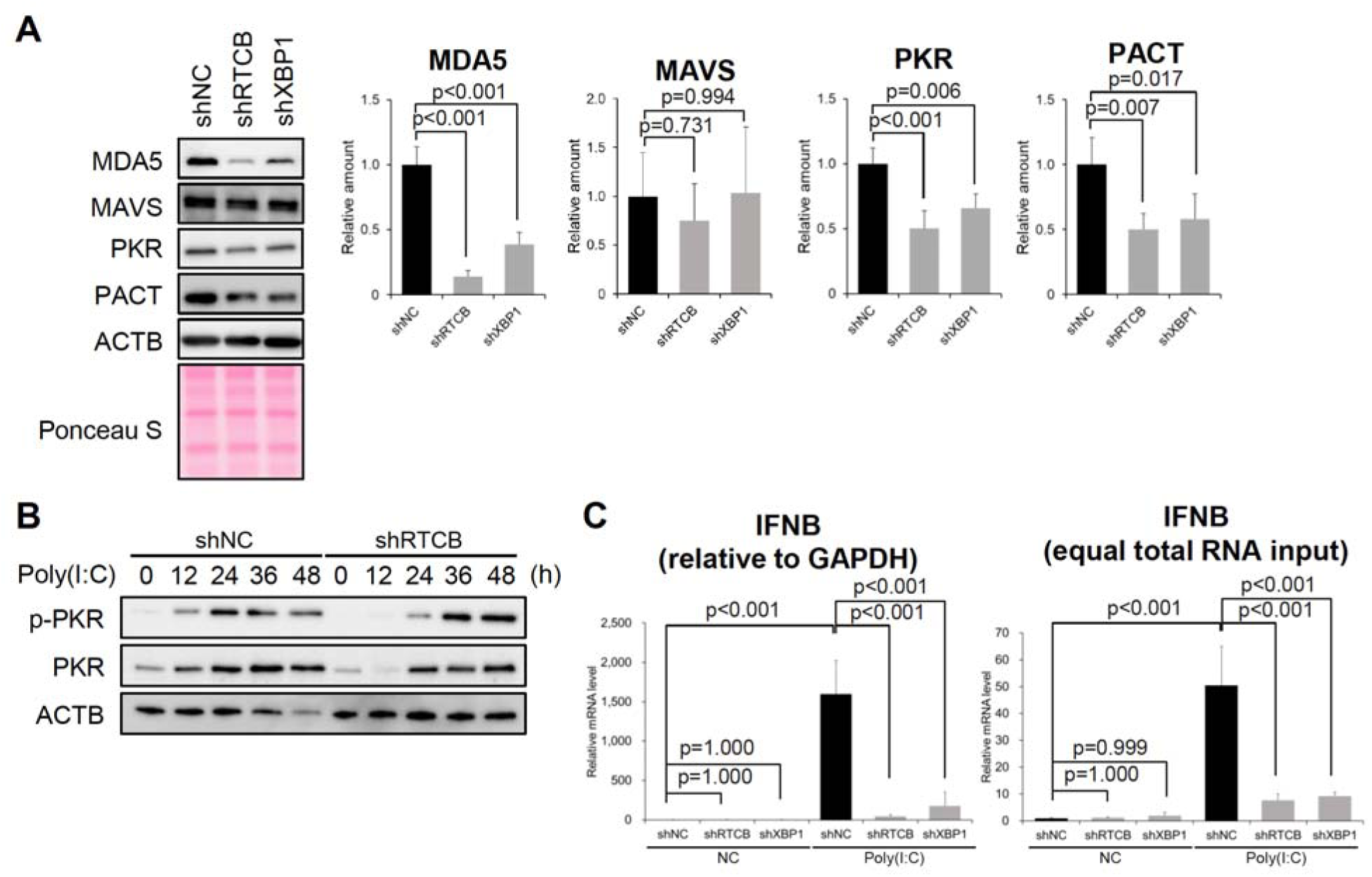
XBP1s positively regulates all three antiviral response pathways. (A) RTCB/XBP1 knockdown reduces the expression of MDA5, PKR and PACT in U2OS cells. Western blotting for MDA5, MAVS, PKR, PACT and ACTB. Relative amounts of MDA5, MAVS, PKR and PACT calculated by densitometry are also shown (n=4 each). (B) RTCB Knockdown delays Poly(I:C)-induced PKR phosphorylation in U2OS cells. Western blotting for phosphorylated PKR (p-PKR), PKR and ACTB. (C) RTCB/XBP1 knockdown decreases Poly(I:C)-induced IFNB mRNA expression in A549 cells. Relative IFNB mRNA levels were determined by ΔΔCT (relative to GAPDH) and equal total RNA input-based quantification. Means and standard deviation were obtained from four independent experiments.

Given that RTCB/XBP1 knockdown decreased PKR and PACT expression in the PKR pathway, we next assessed its impact on PKR pathway activation in U2OS cells (Figure 3B). In control cells (shNC), Poly(I:C)-induced PKR phosphorylation peaked at 24 h and subsequently declined. In contrast, in RTCB knockdown cells (shRTCB), PKR phosphorylation was delayed, peaking at 36 h and remaining at comparable levels even at 48 h (Figure 3B). These findings suggest that XBP1s may modulate the kinetics and magnitude of PKR pathway activation under viral infection.

With respect to the RLR pathway, XBP1s has been reported to enhance transcription of the IFNB gene (29,30). Therefore, reduced production of XBP1s is likely to decrease IFNB gene induction. Because the RTCB complex is involved in XBP1s mRNA splicing (22–24), Poly(I:C)-induced IFNB mRNA induction is expected to be suppressed by knockdown of either RTCB or XBP1. We next examined the impact of RTCB/XBP1 knockdown on RLR pathway activation by measuring IFNB mRNA levels following Poly(I:C) transfection. However, because RNase L is known to cleave a wide range of cytoplasmic RNAs, including housekeeping mRNAs such as GAPDH and ACTB (9,10), normalization to such internal reference genes may compromise the quantitative accuracy of the ΔΔCT method. We therefore measured IFNB gene expression under Poly(I:C) transfection using two quantification approaches: the ΔΔCT method (relative to GAPDH mRNA) and equal total RNA input (Figure 3C). In both quantification methods, knockdown of either RTCB or XBP1 significantly suppressed IFNB mRNA induction (Figure 3C), suggesting that transcription factor XBP1s positively regulates IFNB induction, and that RTCB, acting through the production of XBP1s, also positively regulates IFNB induction. Notably, in the control (shNC), ΔΔCT analysis relative to GAPDH indicated an approximately 1,500-fold induction of IFNB mRNA upon Poly(I:C) transfection, whereas equal total RNA input-based quantification showed an approximately 50-fold increase (Figure 3C). To examine the involvement of RNase L in Poly(I:C)-induced GAPDH mRNA reduction, we compared control and RNase L knockout (ΔRNASEL) A549 cells (Supplemental Fig. S3B, S3C). SYBR Gold staining revealed Poly(I:C)-induced RNA cleavage and tDRs in control cells, whereas these were not detected in ΔRNASEL cells (Supplemental Fig. S3B). Under the condition, GAPDH mRNA levels decreased to approximately one-fortieth after Poly(I:C) transfection, whereas this reduction was completely abolished in ΔRNASEL cells (Supplemental Fig. S3C). These results suggest that the discrepancy between ΔΔCT and equal total RNA input-based quantification can be explained by RNase L-mediated cleavage of GAPDH mRNA. Because the ΔΔCT method may overestimate IFNB induction, we therefore adopted equal total RNA input-based quantification for subsequent experiments. Taken together, these results suggest that transcription factor XBP1s positively regulates the expression of genes constituting the PKR and RLR pathways and further modulates the responsiveness of these pathways. Given that the RNase L pathway was also shown to be regulated by XBP1s (Figure 2), these findings suggest that transcription factor XBP1s controls all three antiviral response pathways triggered by dsRNA.

### RNase L reduces XBP1 mRNA levels and suppresses XBP1s production

A recent study reported that dysfunction of the RNase L pathway underlies a subset of multisystem inflammatory syndrome in children (MIS-C), a rare but severe complication of pediatric SARS-CoV-2 infection characterized by delayed systemic hyperinflammation and multi-organ dysfunction (45). Deficiency of RNase L pathway led to excessive inflammatory cytokine production in response to dsRNA, which was eliminated by MAVS depletion, indicating that the RNase L pathway negatively regulates RLR pathway-dependent inflammation. These data imply that exaggerated RLR pathway-dependent signaling, including enhanced type I interferon responses, may underlie the pathogenesis. However, how the RNase L pathway restrains excessive immune activation via the RLR pathway remains unknown. Given that activated RNase L has been reported to cleave a large proportion of cytoplasmic mRNAs (9,10), and that viral infections, including SARS-CoV-2, lead to endoplasmic reticulum stress due to high levels of viral protein production, resulting in activation of the unfolded protein response (UPR) including XBP1s production (20,46), we hypothesized that RNase L may limit antiviral responses by cleaving XBP1 mRNA and thereby suppressing XBP1s production.

First, we examined whether Poly(I:C)-induced RNase L activation reduces XBP1 mRNA levels in the absence of ER stress (Figure 4A). Consistent with our hypothesis, Poly(I:C) transfection significantly decreased both total XBP1 (XBP1u + XBP1s) mRNA levels and the spliced XBP1s mRNA levels in control cells (Figure 4A). In contrast, no significant decrease was observed in either total XBP1 mRNA or XBP1s mRNA levels in ΔRNASEL cells (Figure 4A). Two-way ANOVA revealed a significant interaction between genotype and Poly(I:C) transfection for both total XBP1 and XBP1s (p<0.001), suggesting that RNase L is required for the Poly(I:C)-induced decrease in XBP1 mRNA expression. We next investigated the effect of RNase L on ER stress-induced XBP1s induction. To evaluate ER stress response associated with viral infection, we employed a combination of Poly(I:C) transfection and thapsigargin treatment. At 4 hours after thapsigargin treatment, XBP1s mRNA levels were significantly increased in both control and ΔRNASEL cells (Figure 4B), with no significant interaction between genotype and Poly(I:C) treatment (two-way ANOVA, p=0.688). At 8 hours after stimulation in the Poly(I:C)-thapsigargin combination model, Poly(I:C) transfection significantly decreased XBP1s mRNA levels, and this decrease was abolished in ΔRNASEL cells (Figure 4C), suggesting that active RNase L suppresses ER stress-mediated XBP1s mRNA induction. We also examined the effect of RNase L activation on XBP1s mRNA levels under ER stress conditions using a time-course analysis. Compared with control cells, ΔRNASEL cells exhibited markedly elevated XBP1s mRNA levels at 4 and 8 hours, whereas by 24 hours the levels declined to those comparable to controls (Figure 4D). In contrast, XBP1u pre-mRNA levels decreased similarly in both control and ΔRNASEL cells at 4 hours. By 8 hours, however, XBP1u expression increased in ΔRNASEL cells, and at 24 hours remained elevated relative to control cells (Supplemental Fig. S4A), indicating that RNase L negatively regulates not only XBP1s mRNA but also XBP1u pre-mRNA levels in the Poly(I:C)-thapsigargin combination model.

**Figure 4:**
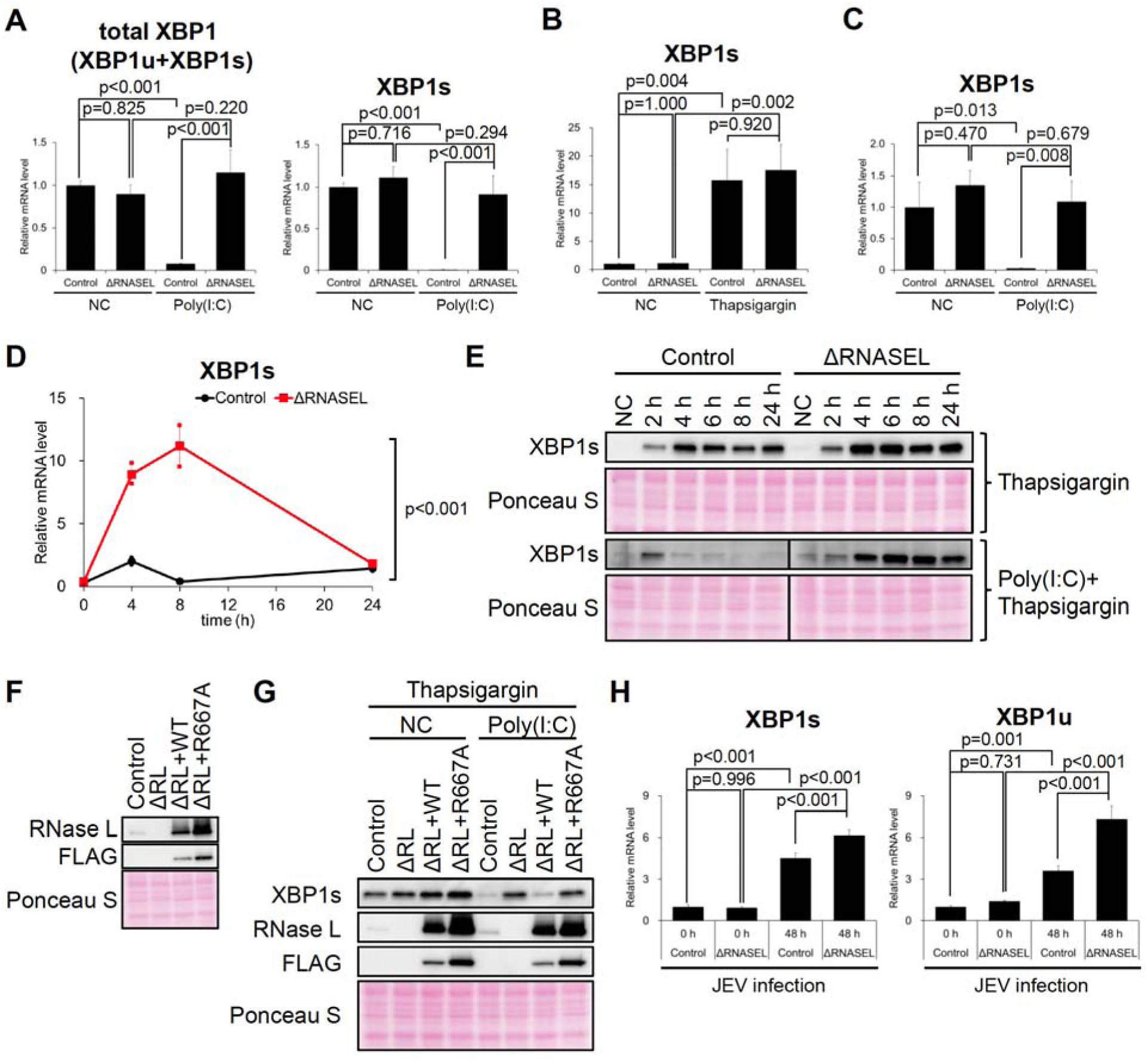
RNase L reduces XBP1 mRNA levels and suppresses XBP1s production. (A) Active RNase L reduces XBP1s mRNA levels in A549 cells. Relative expression levels of total XBP1 (XBP1u and XBP1s) and XBP1s mRNA are shown. (B) Thapsigargin treatment for 4 hours induced comparable levels of XBP1s mRNA in control and ΔRNASEL cells. (C) Active RNase L also decreases XBP1s mRNA levels in the Poly(I:C)-thapsigargin combination model. Relative XBP1s mRNA expression was quantified by qPCR 8 hours after treatment with Poly(I:C) and thapsigargin. (D) Time-dependent induction of XBP1s mRNA in the Poly(I:C)-thapsigargin combination model. (E) Time-dependent induction of transcription factor XBP1s in the Poly(I:C)-thapsigargin combination model. Western blot analysis of XBP1s is shown. (F, G) RNase activity of RNase L is required for suppression of XBP1s induction. (F) Western blot analysis of RNase L and FLAG. ΔRL, RNase L knockout (ΔRNASEL); ΔRL + WT, ΔRL cells reconstituted with wild-type RNase L; ΔRL + R667A, ΔRL cells reconstituted with an inactive RNase L mutant. (G) RNase L reconstitution restored suppression of XBP1s induction in ΔRNASEL cells. XBP1s protein levels were evaluated by Western blotting 6 hours after treatment with 25 nM thapsigargin and 2 µg/mL Poly(I:C). (H) RNase L significantly represses XBP1 mRNA induction during JEV infection. Relative expression levels of XBP1s mRNA and XBP1u pre-mRNA quantified based on equal total RNA input. Data in (A)-(D) and (H) are from three independent experiments.

Given the observed changes at the mRNA level, we next assessed the impact of RNase L on XBP1s protein by time-course Western blotting in the Poly(I:C)-thapsigargin combination model (Figure 4E). With thapsigargin treatment alone, XBP1s was induced from 2 hours in both control and ΔRNASEL cells, peaked at 4 hours, and remained at comparable levels up to 24 hours. In contrast, under the Poly(I:C)-thapsigargin combination condition, XBP1s was induced at 2 hours in control cells but was suppressed at 4 hours, and this suppression persisted up to 24 hours. In ΔRNASEL cells, the suppression of XBP1s observed in control cells was not detected, and the expression pattern resembled that observed with thapsigargin alone (Figure 4E). These results are not inconsistent with the time-course pattern of XBP1s mRNA levels described above (Figure 4D). Given that RNase L is activated as early as 2 hours following Poly(I:C) transfection (10), the behavior of XBP1s mRNA and protein levels observed here may be attributable to the rapid activation of RNase L. To determine whether the Poly(I:C)-induced XBP1s reduction depends on the enzymatic activity of RNase L, we generated ΔRNASEL cells reconstituted with either wild-type RNASEL or an enzymatically inactive mutant (R667A) RNASEL (Figure 4F). Initially, we examined whether reconstitution of wild-type RNASEL in ΔRNASEL cells restored the enzymatic activity of RNase L. As shown in Supplemental Fig. S4B and S4C, loss of Poly(I:C)-induced tDR production, including 5’-tDR^Ala^, in ΔRNASEL cells was rescued by wild-type RNase L knock-in (ΔRL+WT) but not by inactive RNase L (ΔRL+R667A). The levels of both tDRs and 5′-tDR^Ala^ were higher in ΔRL+WT cells than in control cells (Supplemental Fig. S4B, C), likely due to the higher RNase L levels in ΔRL+WT cells (Figure 4F). Under the condition, in ΔRL+WT cells, XBP1s was reduced upon Poly(I:C) transfection, similar to control cells, whereas ΔRL+R667A cells did not exhibit this reduction and showed XBP1s levels comparable to those in ΔRNASEL cells (Figure 4G). These results demonstrate that the reduction of Poly(I:C)-mediated XBP1s levels is dependent on the enzymatic activity of RNase L, thereby further supporting the idea that RNase L reduces the levels of XBP1s transcription factor through cleavage of XBP1 mRNA.

We further investigated the impact of RNase L on XBP1 mRNA expression in a Japanese encephalitis virus (JEV) infection model, as JEV, a member of the Flaviviridae family, has been reported to trigger the UPR (47–49). At 48 hours post-infection, both control and ΔRNASEL cells exhibited significant increases in XBP1s and XBP1u mRNA levels. However, expression levels at 48 hours were significantly higher in ΔRNASEL cells (Figure 4H, Supplemental Fig. S4D). Two-way ANOVA revealed significant interactions between genotype and time for both XBP1s and XBP1u (p=0.001 and p<0.001 in Figure 4H, respectively), suggesting that RNase L contributes to the regulation of XBP1 mRNA levels during JEV infection.

Taken together, the in vitro and JEV infection data indicate that RNase L functions as a negative regulator of XBP1s abundance during viral infection, likely by cleaving XBP1 mRNA via its RNase activity.

### RNase L limits IFNB mRNA induction by reducing XBP1s production

We further investigated the relationship between XBP1s and the RLR pathway activation observed in ΔRNASEL cells under the Poly(I:C)-thapsigargin combination condition, by measuring IFNB mRNA levels, a transcriptional target induced upon the activation of RLR pathway. At 24 hours after Poly(I:C) transfection and thapsigargin treatment, IFNB mRNA levels were significantly increased in both control and ΔRNASEL cells. However, expression levels were significantly higher in ΔRNASEL cells than in control cells (Figure 5A, Supplemental Fig. S5A), consistent with a previous report (45).

**Figure 5:**
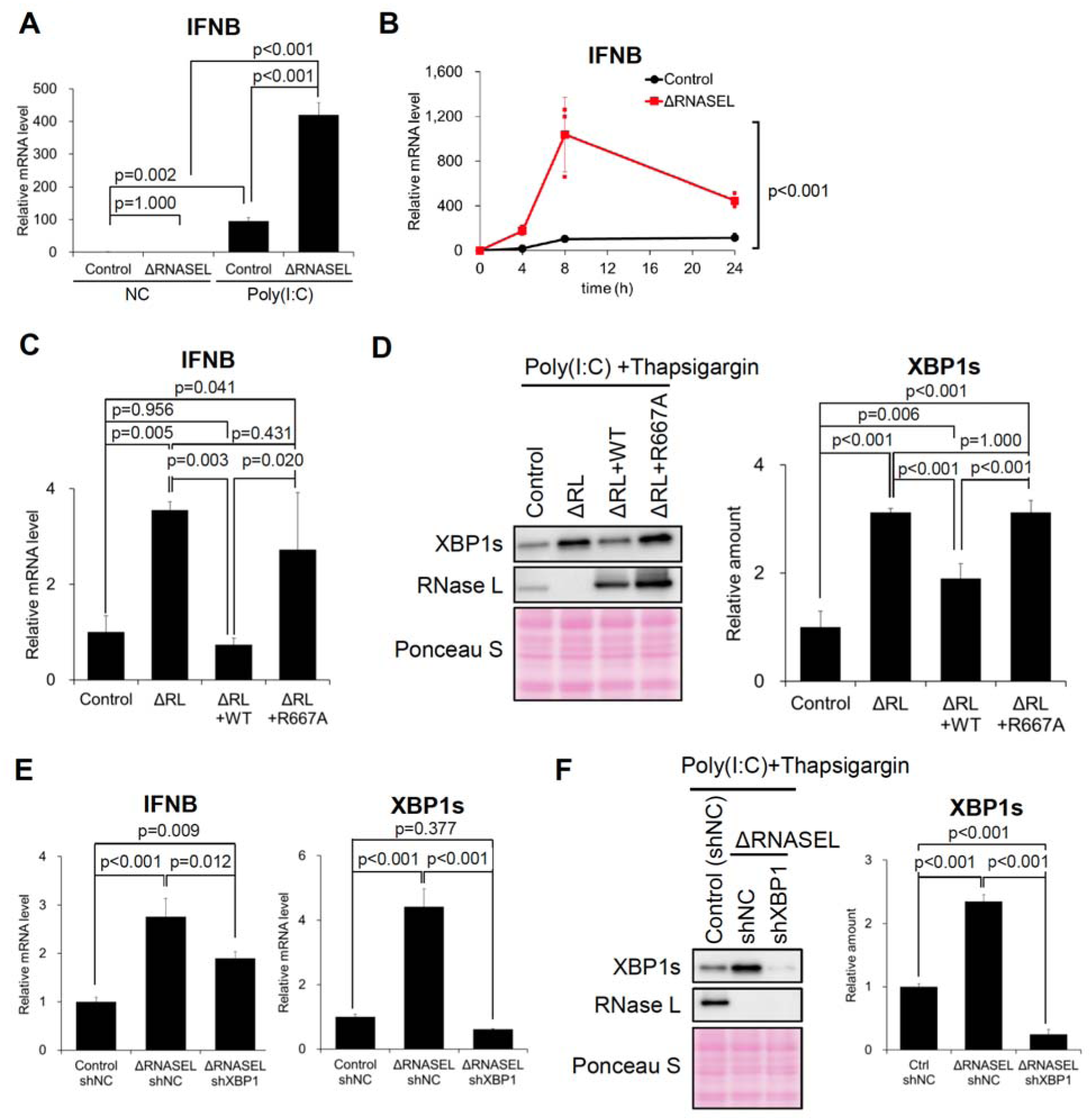
RNase L limits IFNB mRNA induction by reducing XBP1s production. (A, B) RNase L deletion enhances IFNB mRNA induction in the Poly(I:C)-thapsigargin combination model. (A) IFNB mRNA levels 24 hours after treatment in control and ΔRNASEL cells. (B) Time course of IFNB mRNA induction. (C, D) RNase activity of RNase L is required for suppression of Poly(I:C)-mediated IFNB mRNA induction. (C) Relative expression levels of IFNB mRNA and (D) Western blotting for XBP1s and RNase L. IFNB mRNA and XBP1s levels were evaluated at 8 hours in the Poly(I:C)-thapsigargin combination model. ΔRL, RNase L knockout (ΔRNASEL); ΔRL + WT, ΔRL cells reconstituted with wild-type RNase L; ΔRL + R667A, ΔRL cells reconstituted with an inactive RNase L mutant. (E, F) XBP1 knockdown suppresses enhanced IFNB mRNA induction in ΔRNASEL cells in the Poly(I:C)-thapsigargin combination model. (E) Relative expression levels of IFNB and XBP1s mRNA and (D) Western blotting for XBP1s and RNase L at 8 hours in the Poly(I:C)-thapsigargin combination model. Data were obtained from three independent experiments.

In the time-course experiment, equal total RNA input-based quantification revealed elevated IFNB mRNA levels in ΔRNASEL cells relative to control cells between 4 and 24 hours (Figure 5B), whereas ΔΔCt analysis showed comparable behavior in the two genotypes up to 8 hours, with increased IFNB mRNA expression in ΔRNASEL cells becoming apparent only at 24 hours (Supplemental Fig. S5B). Two-way ANOVA revealed significant genotype-time interactions in both equal total RNA input-based quantification and ΔΔCt analysis (p < 0.001), suggesting that RNase L is involved in regulating IFNB expression in the Poly(I:C)-thapsigargin combination model. We next examined whether the enzymatic activity of RNase L is required to suppress the enhanced induction of IFNB mRNA in ΔRNASEL cells (Figure 5C, D). The increase in IFNB mRNA observed in ΔRNASEL cells was completely abolished by knock-in of wild-type RNase L (ΔRL+WT), returning to control levels, whereas knock-in of enzymatically inactive RNase L (ΔRL+R667A) failed to reduce IFNB mRNA levels (Figure 5C). Under this condition, XBP1s protein levels exhibited a pattern parallel to that of IFNB mRNA, with comparable responses to wild-type and inactive RNase L knock-in (Figure 5D). These data suggest that the suppression of IFNB mRNA by RNase L is dependent on its enzymatic activity, likely via XBP1 mRNA cleavage. Finally, we examined whether the exaggerated induction of IFNB mRNA observed in ΔRNASEL cells could be suppressed by XBP1 knockdown (Figure 5E, F). In the Poly(I:C)-thapsigargin combination model, XBP1 knockdown significantly reduced IFNB mRNA levels in ΔRNASEL cells (Figure 5E). Under the same condition, both XBP1s mRNA (Figure 5E) and XBP1s protein levels (Figure 5F) were significantly decreased by XBP1 knockdown. These findings indicate that the exaggerated induction of IFNB mRNA caused by RNASEL knockout can be attenuated by XBP1 knockdown.

## DISCUSSION

In this study, we demonstrated that XBP1s enhances all three major antiviral response pathways. We also revealed that, RNase L, an effector of the antiviral response, suppresses XBP1s production by cleaving XBP1 mRNA, thereby potentially preventing excessive XBP1s-driven immune responses. Our findings reveal a previously unrecognized mechanism of antiviral response regulation by RNase L and thus provide a framework for understanding how antiviral signaling is fine-tuned during cellular stress.

Whereas XBP1s has been reported to promote IFNB transcription in the RLR pathway (29,30), whether XBP1s also regulates the responsiveness of the RNase L and PKR pathways has remained unclear. In the present study, we revealed that RTCB/XBP1 knockdown not only suppressed IFNB mRNA induction (Figure 3C) but also reduced RNase L-mediated tDR production (Figure 2H, I) and delayed PKR phosphorylation (Figure 3B). These findings suggest that XBP1s positively regulates the responsiveness of all three antiviral response pathways and that RTCB may also participate in this regulation through control of XBP1s production. Although we were not able to confirm changes in all genes constituting the three antiviral response pathways, RTCB/XBP1 knockdown reduced the expression of RNASEL, OASs, MDA5, PKR and PACT (Figure 2C, 3A, Supplemental Fig. S2B, S3A), raising the possibility that this reduction contributed to attenuation of responsiveness in each pathway. It is important to note that, in addition to our finding that XBP1s positively regulates MDA5 expression, XBP1s directly enhances IFNB transcription in the RLR pathway (29,30). Under the assumption that XBP1s is generated during viral infection, changes in the expression of dsRNA sensors and effector molecules may require additional time to affect pathway responsiveness. In contrast, direct enhancement of IFNB mRNA transcription is more immediately connected to subsequent interferon signaling. These considerations raise the possibility that XBP1s is relatively more important for regulation of the RLR pathway during the acute phase of infection.

Our results demonstrated that activation of RNase L rapidly reduces XBP1 mRNA and XBP1s protein levels (Figure 4D, E). In addition, rescue experiments in which wild-type or inactive RNase L was reintroduced into ΔRNASEL cells showed that the RNase L-dependent suppression of XBP1 expression requires its RNase activity (Figure 4F, G), raising the possibility that this effect is mediated by cleavage of XBP1 mRNA. Notably, XBP1s has been reported to be a short-lived protein with a half-life of approximately 10 min under basal conditions (50,51), and its production is tightly coupled to ER stress through the inducible splicing of XBP1u mRNA. XBP1u mRNA is a unique transcript that is cotranslationally recruited to the ER membrane (52) and, in response to ER stress, undergoes IRE1α-mediated cleavage and RTCB complex-dependent ligation to generate XBP1s mRNA (22–24), which is then translated into the transcription factor XBP1s. Thus, stress-induced XBP1s production can proceed from pre-existing XBP1u mRNA without an immediate requirement for de novo XBP1 transcription, enabling a rapid response to ER stress. These unique features may allow RNase L-mediated cleavage of XBP1 mRNA to rapidly attenuate XBP1s expression.

Previous studies have described mechanisms by which RNase L enhances antiviral responses, particularly IFN production (14,53), as well as evidence suggesting that RNase L can also function suppressively in antiviral responses (54–56), suggesting that its effects are context-dependent. Our findings extend these observations by suggesting a new molecular mechanism through which RNase L negatively regulates antiviral responses. Interestingly, Banerjee et al. reported that, in murine macrophages, RNase L suppresses IFN-β induction and that this suppressive effect depends on the high activity of the RNase L/OAS pathway in these cells (56). This observation is consistent with our findings. Lee et al. further reported, based on analyses of MIS-C cases, that the OAS-RNase L pathway suppresses RLR-MAVS-mediated inflammatory cytokine production (45). However, the molecular mechanism by which RNase L restrains such cytokine responses was not defined in that study. Our findings raise the possibility that RNase L-mediated cleavage of XBP1 mRNA, leading to reduced XBP1s production, may represent one mechanism underlying this suppressive effect. SARS-CoV-2 infection has been reported to perturb ER homeostasis and activate the unfolded protein response (57,58), although the extent of activation of the IRE1α-XBP1 branch appears to vary depending on the experimental context. Thus, the RNase L-dependent regulation of XBP1s identified here may contribute, at least in part, to the pathogenesis of MIS-C. More broadly, as XBP1s is widely known to potentiate immune responses across both innate and adaptive immunity (26), the rapid suppression of XBP1s expression by RNase L identified here may have broader consequences for immune responsiveness beyond antiviral signaling alone.

One particularly intriguing implication of our findings is that RNase L, an effector of one branch of the antiviral response, appears to form a negative-feedback loop that limits the XBP1s-mediated enhancement of antiviral responsiveness. Mechanistically, this putative feedback loop would involve at least three virus-triggered processes: (i) XBP1s induction as part of the ER stress response, (ii) activation of the RLR pathway, and (iii) activation of RNase L. For XBP1s to enhance antiviral responsiveness, particularly IFN induction through the RLR pathway, RLR signaling would need to be activated and XBP1s to be induced by the UPR before RNase L activation reaches a sufficient level. Although recent studies have shown that viral RNA sensing by RIG-I and activation of the RLR pathway can occur very rapidly (59), the relative kinetics of the antiviral branches examined here, especially the RLR and OAS-RNase L pathways, as well as the temporal order of XBP1s induction and antiviral pathway activation during viral infection, remain largely undefined and are likely to vary depending on the virus and the infected cell type. Our data therefore support a model in which, once activated, RNase L rapidly suppresses further XBP1s induction by cleaving XBP1 mRNA, thereby limiting XBP1s-dependent amplification of antiviral signaling; however, the extent to which this mechanism contributes to antiviral control is also likely to be virus- and cell-context dependent. Further studies will be needed to identify the types of viral infections and tissues in which this mechanism plays a major role. Importantly, our findings also suggest that, during viral infection, RNase L may regulate not only antiviral responses but also XBP1s-mediated UPR signaling. It will therefore be important to investigate how RNase L-dependent suppression of XBP1s affects ER homeostasis under conditions of viral infection, and whether this effect is ultimately beneficial or detrimental to the host.

Transfer RNA-derived RNAs (tDRs), especially those induced under stress conditions, have emerged as functional stress-response molecules generated by the cleavage of mature tRNAs or precursor tRNAs (pre-tRNAs) by RNases, including ANG, and are thought to contribute to cellular stress adaptation through mechanisms such as translational repression (17,34). We have previously shown that the RTCB-LC negatively regulates ANG-induced tiRNA production by ligating cleaved tiRNAs, and that tiRNA production is further increased under oxidative stress through inhibition of RTCB (35). In the present study, we found that the RTCB-LC regulates the production of RNase L-induced tDRs not only through direct repair of cleaved tRNAs as in the case of ANG-induced tiRNAs, but also by positively regulating the extent of RNase L-mediated tRNA cleavage through regulation of XBP1s production. Because RNase L-induced tDRs include 5’-tDR^Ala^, which has translational repressive activity like ANG-induced 5’-tiRNA^Ala^ (18), RTCB-dependent regulation of RNase L-induced tDR production may affect stress adaptation during viral infection. Given that viral infection can induce oxidative stress (60), oxidative inhibition of the RTCB complex could amplify the production of RNase L-induced tDRs, which in turn may contribute to cytoprotective effects in infected cells. In that case, what is likely to happen to tDR production if the RTCB-LC is suppressed by oxidative stress during viral infection? Because rapid inhibition of RTCB by H_2_O_2_ significantly increased RNase L-induced tDR levels (Figure 1C, D, Supplemental Fig. S1), it is plausible that, at an early stage of infection, reduced RTCB-dependent tRNA repair has a greater impact, leading to increased tDR production. This may enhance the cytoprotective effects of RNase L-induced tDRs during the acute phase. In contrast, when RTCB suppression is prolonged, reduced XBP1s production may lower the expression of RNase L and OASs, attenuate RNase L-mediated tRNA cleavage, and thereby decrease tDR production (Figure 1E-G, Supplemental Fig. S2B-D). Thus, if viral infection persists and RTCB activity remains suppressed over time, tDR production may eventually decline because RNase L responsiveness becomes attenuated. Because host responses to viral infection, including the UPR, antiviral responses, and oxidative stress, are highly complex, further studies will be needed to clarify how this regulation of RNase L-induced tDRs contributes to cellular stress responses during viral infection.

In summary, our study demonstrated that the transcription factor XBP1s broadly enhances antiviral responsiveness and further reveals a negative-feedback mechanism whereby RNase L, an effector of one branch of the antiviral response, rapidly suppresses XBP1s production through cleavage of XBP1 mRNA. Beyond the enhancement of antiviral responsiveness identified here, XBP1s has also been reported to potentiate both innate and adaptive immunity, and is essential for the maintenance of ER homeostasis during ER stress. Therefore, the rapid regulation of XBP1s by RNase L identified in this study is likely to have broad consequences for cellular stress responses during viral infection. Further elucidation of this regulatory axis may provide new insight into how host stress responses are coordinated during viral infection.

## ACKNOWLEDGEMENTS

AUTHOR CONTRIBUTIONS

Y. Takenaka (Conceptualization [equal], Formal Analysis [lead], Funding acquisition [supporting], Investigation [lead], Methodology [equal], Validation [lead], Visualization [lead], Writing –original draft [equal], Writing –review & editing [equal]), Y.A. (Conceptualization [equal], Formal Analysis [equal], Funding acquisition [supporting], Investigation [equal], Methodology [equal], Project administration [lead], Supervision [equal], Visualization [equal], Writing –original draft [equal], Writing –review & editing [equal]), T.I. (Investigation [supporting], Resources [supporting]), D.S. (Investigation [supporting], Validation [supporting]), K.A. (Resources [supporting]), R.O. (Investigation [supporting], Resources [supporting]), N.K. (Resources [supporting]), T.A. (Funding acquisition [equal], Project administration [equal], Supervision [equal]), E.M. (Funding acquisition [supporting], Project administration [equal], Supervision [equal], Writing –original draft [equal], Writing –review & editing [equal]), Y. Tomioka (Funding acquisition [equal], Project administration [equal], Supervision [equal]), and P.I. (Conceptualization [equal], Funding acquisition [equal], Project administration [equal], Supervision [equal], Writing –original draft [equal], Writing –review & editing [equal])

## SUPPLEMENTARY DATA

Supplementary Data are available online.

## FUNDING

This work was supported by NIH [R01GM146997 to P.I.], JSPS KAKENHI [23KK0290 to Y.A., 23K06736 to Y. Tomioka, 23K24135 to E.M.], the Japan Agency for Medical Research and Development (AMED) [24zf0127001h0004 to T.A.], and JST SPRING [JPMJSP2114 to Y.

Takenaka].

## DATA AVAILABILITY

The data underlying this article will be shared on reasonable request to the corresponding author.

APPENDIX

## REFERENCES

1. Wang, R., Lan, C., Benlagha, K., Camara, N.O.S., Miller, H., Kubo, M., Heegaard, S., Lee, P., Yang, L., Forsman, H., et al. (2024) The interaction of innate immune and adaptive immune system. MedComm (2020), 5, e714.

2. Carroll, S.L., Pasare, C. and Barton, G.M. (2024) Control of adaptive immunity by pattern recognition receptors. Immunity, 57, 632–648.

3. Hur, S. (2019) Double-Stranded RNA Sensors and Modulators in Innate Immunity. Annu Rev Immunol, 37, 349–375.

4. Cottrell, K.A., Andrews, R.J. and Bass, B.L. (2024) The competitive landscape of the dsRNA world. Mol Cell, 84, 107–119.

5. Pakos-Zebrucka, K., Koryga, I., Mnich, K., Ljujic, M., Samali, A. and Gorman, A.M. (2016) The integrated stress response. EMBO Rep, 17, 1374–1395.

6. Costa-Mattioli, M. and Walter, P. (2020) The integrated stress response: From mechanism to disease. Science, 368, aat5314.

7. Schwartz, S.L. and Conn, G.L. (2019) RNA regulation of the antiviral protein 2’-5’-oligoadenylate synthetase. Wiley Interdiscip Rev RNA, 10, e1534.

8. Karasik, A. and Guydosh, N.R. (2024) The Unusual Role of Ribonuclease L in Innate Immunity. Wiley Interdiscip Rev RNA, 15, e1878.

9. Rath, S., Prangley, E., Donovan, J., Demarest, K., Wingreen, N.S., Meir, Y. and Korennykh, A. (2019) Concerted 2-5A-Mediated mRNA Decay and Transcription Reprogram Protein Synthesis in the dsRNA Response. Mol Cell, 75, 1218–1228 e1216.

10. Burke, J.M., Moon, S.L., Matheny, T. and Parker, R. (2019) RNase L Reprograms Translation by Widespread mRNA Turnover Escaped by Antiviral mRNAs. Mol Cell, 75, 1203–1217 e1205.

11. Xi, J., Snieckute, G., Martinez, J.F., Arendrup, F.S.W., Asthana, A., Gaughan, C., Lund, A.H., Bekker-Jensen, S. and Silverman, R.H. (2024) Initiation of a ZAKalpha-dependent ribotoxic stress response by the innate immunity endoribonuclease RNase L. Cell Rep, 43, 113998.

12. Karasik, A., Lorenzi, H.A., DePass, A.V. and Guydosh, N.R. (2024) Endonucleolytic RNA cleavage drives changes in gene expression during the innate immune response. Cell Rep, 43, 114287.

13. Burke, J.M., Gilchrist, A.R., Sawyer, S.L. and Parker, R. (2021) RNase L limits host and viral protein synthesis via inhibition of mRNA export. Sci Adv, 7, abh2479.

14. Malathi, K., Dong, B., Gale, M., Jr. and Silverman, R.H. (2007) Small self-RNA generated by RNase L amplifies antiviral innate immunity. Nature, 448, 816–819.

15. Donovan, J., Rath, S., Kolet-Mandrikov, D. and Korennykh, A. (2017) Rapid RNase L-driven arrest of protein synthesis in the dsRNA response without degradation of translation machinery. RNA, 23, 1660–1671.

16. Holmes, A.D., Chan, P.P., Chen, Q., Ivanov, P., Drouard, L., Polacek, N., Kay, M.A. and Lowe, T.M. (2023) A standardized ontology for naming tRNA-derived RNAs based on molecular origin. Nat Methods, 20, 627–628.

17. Akiyama, Y. and Ivanov, P. (2023) tRNA-derived RNAs: Biogenesis and roles in translational control. *Wiley Interdiscip Rev RNA*, e1805.

18. Takenaka, Y., Yamada, A., Tomioka, Y., Akiyama, Y. and Ivanov, P. (2025) RNase L produces tRNA-derived RNAs that contribute to translation inhibition. RNA, 31, 961–972.

19. Prasad, V. and Greber, U.F. (2021) The endoplasmic reticulum unfolded protein response - homeostasis, cell death and evolution in virus infections. FEMS Microbiol Rev, 45.

20. Chen, H., Liu, D., Hua, J., Wu, M., Hua, Y., Feng, C., He, Z., Moffett, P. and Zhang, K. (2026) Hijacking the unfolded protein response (UPR) pathway: Balancing viral infection and host cell survival. The Journal of biological chemistry, 302, 111040.

21. Hetz, C., Zhang, K. and Kaufman, R.J. (2020) Mechanisms, regulation and functions of the unfolded protein response. Nat Rev Mol Cell Biol, 21, 421–438.

22. Lu, Y., Liang, F.X. and Wang, X. (2014) A synthetic biology approach identifies the mammalian UPR RNA ligase RtcB. Mol Cell, 55, 758–770.

23. Kosmaczewski, S.G., Edwards, T.J., Han, S.M., Eckwahl, M.J., Meyer, B.I., Peach, S., Hesselberth, J.R., Wolin, S.L. and Hammarlund, M. (2014) The RtcB RNA ligase is an essential component of the metazoan unfolded protein response. EMBO Rep, 15, 1278–1285.

24. Jurkin, J., Henkel, T., Nielsen, A.F., Minnich, M., Popow, J., Kaufmann, T., Heindl, K., Hoffmann, T., Busslinger, M. and Martinez, J. (2014) The mammalian tRNA ligase complex mediates splicing of XBP1 mRNA and controls antibody secretion in plasma cells. The EMBO journal, 33, 2922–2936.

25. Park, S.M., Kang, T.I. and So, J.S. (2021) Roles of XBP1s in Transcriptional Regulation of Target Genes. Biomedicines, 9.

26. Di Conza, G., Ho, P.C., Cubillos-Ruiz, J.R. and Huang, S.C. (2023) Control of immune cell function by the unfolded protein response. Nat Rev Immunol, 23, 546–562.

27. Martinon, F., Chen, X., Lee, A.H. and Glimcher, L.H. (2010) TLR activation of the transcription factor XBP1 regulates innate immune responses in macrophages. Nat Immunol, 11, 411–418.

28. Iwakoshi, N.N., Pypaert, M. and Glimcher, L.H. (2007) The transcription factor XBP-1 is essential for the development and survival of dendritic cells. J Exp Med, 204, 2267–2275.

29. Zeng, L., Liu, Y.P., Sha, H., Chen, H., Qi, L. and Smith, J.A. (2010) XBP-1 couples endoplasmic reticulum stress to augmented IFN-beta induction via a cis-acting enhancer in macrophages. J Immunol, 185, 2324–2330.

30. Smith, J.A., Turner, M.J., DeLay, M.L., Klenk, E.I., Sowders, D.P. and Colbert, R.A. (2008) Endoplasmic reticulum stress and the unfolded protein response are linked to synergistic IFN-beta induction via X-box binding protein 1. Eur J Immunol, 38, 1194–1203.

31. Popow, J., Englert, M., Weitzer, S., Schleiffer, A., Mierzwa, B., Mechtler, K., Trowitzsch, S., Will, C.L., Luhrmann, R., Soll, D. et al. (2011) HSPC117 is the essential subunit of a human tRNA splicing ligase complex. Science, 331, 760–764.

32. Lyons, S.M., Fay, M.M., Akiyama, Y., Anderson, P.J. and Ivanov, P. (2017) RNA biology of angiogenin: Current state and perspectives. RNA Biol, 14, 171–178.

33. Ivanov, P., Emara, M.M., Villen, J., Gygi, S.P. and Anderson, P. (2011) Angiogenin-induced tRNA fragments inhibit translation initiation. Mol Cell, 43, 613–623.

34. Akiyama, Y. and Ivanov, P. (2024) Oxidative Stress, Transfer RNA Metabolism, and Protein Synthesis. Antioxid Redox Signal, 40, 715–735.

35. Akiyama, Y., Takenaka, Y., Kasahara, T., Abe, T., Tomioka, Y. and Ivanov, P. (2022) RTCB Complex Regulates Stress-Induced tRNA Cleavage. Int J Mol Sci, 23, 13100.

36. Asanovic, I., Strandback, E., Kroupova, A., Pasajlic, D., Meinhart, A., Tsung-Pin, P., Djokovic, N., Anrather, D., Schuetz, T., Suskiewicz, M.J. et al. (2021) The oxidoreductase PYROXD1 uses NAD(P)(+) as an antioxidant to sustain tRNA ligase activity in pre-tRNA splicing and unfolded protein response. Mol Cell, 81, 2520–2532 e2516.

37. Tanaka, N., Meineke, B. and Shuman, S. (2011) RtcB, a novel RNA ligase, can catalyze tRNA splicing and HAC1 mRNA splicing in vivo. The Journal of biological chemistry, 286, 30253–30257.

38. Chakravarty, A.K., Subbotin, R., Chait, B.T. and Shuman, S. (2012) RNA ligase RtcB splices 3’-phosphate and 5’-OH ends via covalent RtcB-(histidinyl)-GMP and polynucleotide-(3’)pp(5’)G intermediates. Proceedings of the National Academy of Sciences of the United States of America, 109, 6072–6077.

39. Naito, Y., Yoshimura, J., Morishita, S. and Ui-Tei, K. (2009) siDirect 2.0: updated software for designing functional siRNA with reduced seed-dependent off-target effect. BMC Bioinformatics, 10, 392.

40. Takenaka, Y., Aoyama, K. and Akiyama, Y. (2025) Northern blotting for human pre-tRNA and tRNA-derived RNAs. Methods Enzymol, 711, 15–27.

41. Ii, K., Sato, F., Hatakeyama, Y., Suzuki, H., Noguchi, T., Ishida, K., Arakawa, M., Nakamura, K., Iwatsuki, R., Nguyen, C.T. et al. (2025) Flavivirus-based bivalent nanoparticle vaccines induce neutralizing antibodies and Th1 responses against flavivirus and coupling antigens. iScience, 28, 113659.

42. Yang, W. (2011) Nucleases: diversity of structure, function and mechanism. Q Rev Biophys, 44, 1–93.

43. Li, Y., Banerjee, S., Wang, Y., Goldstein, S.A., Dong, B., Gaughan, C., Silverman, R.H. and Weiss, S.R. (2016) Activation of RNase L is dependent on OAS3 expression during infection with diverse human viruses. Proceedings of the National Academy of Sciences of the United States of America, 113, 2241–2246.

44. Chukwurah, E., Farabaugh, K.T., Guan, B.J., Ramakrishnan, P. and Hatzoglou, M. (2021) A tale of two proteins: PACT and PKR and their roles in inflammation. FEBS J, 288, 6365–6391.

45. Lee, D., Le Pen, J., Yatim, A., Dong, B., Aquino, Y., Ogishi, M., Pescarmona, R., Talouarn, E., Rinchai, D., Zhang, P., et al. (2023) Inborn errors of OAS-RNase L in SARS-CoV-2-related multisystem inflammatory syndrome in children. Science, 379, eabo3627.

46. Keramidas, P., Pitou, M., Papachristou, E. and Choli-Papadopoulou, T. (2024) Insights into the Activation of Unfolded Protein Response Mechanism during Coronavirus Infection. Curr Issues Mol Biol, 46, 4286–4308.

47. Verhaegen, M. and Vermeire, K. (2024) The endoplasmic reticulum (ER): a crucial cellular hub in flavivirus infection and potential target site for antiviral interventions. Npj Viruses, 2, 24.

48. Su, H.L., Liao, C.L. and Lin, Y.L. (2002) Japanese encephalitis virus infection initiates endoplasmic reticulum stress and an unfolded protein response. Journal of virology, 76, 4162–4171.

49. Kumar, S., Verma, A., Yadav, P., Dubey, S.K., Azhar, E.I., Maitra, S.S. and Dwivedi, V.D. (2022) Molecular pathogenesis of Japanese encephalitis and possible therapeutic strategies. Arch Virol, 167, 1739–1762.

50. Sun, H., Wei, G., Liu, H., Xiao, D., Huang, J., Lu, J., Miao, J., Liu, J. and Chen, S. (2020) Inhibition of XBP1s ubiquitination enhances its protein stability and improves glucose homeostasis. Metabolism: clinical and experimental, 105, 154046.

51. Liu, J., Ibi, D., Taniguchi, K., Lee, J., Herrema, H., Akosman, B., Mucka, P., Salazar Hernandez, M.A., Uyar, M.F., Park, S.W. et al. (2016) Inflammation Improves Glucose Homeostasis through IKKbeta-XBP1s Interaction. Cell, 167, 1052–1066 e1018.

52. Yanagitani, K., Kimata, Y., Kadokura, H. and Kohno, K. (2011) Translational pausing ensures membrane targeting and cytoplasmic splicing of XBP1u mRNA. Science, 331, 586–589.

53. Manivannan, P., Siddiqui, M.A. and Malathi, K. (2020) RNase L Amplifies Interferon Signaling by Inducing Protein Kinase R-Mediated Antiviral Stress Granules. Journal of virology, 94.

54. Li, X.L., Blackford, J.A., Judge, C.S., Liu, M., Xiao, W., Kalvakolanu, D.V. and Hassel, B.A. (2000) RNase-L-dependent destabilization of interferon-induced mRNAs. A role for the 2-5A system in attenuation of the interferon response. The Journal of biological chemistry, 275, 8880–8888.

55. Khabar, K.S., Siddiqui, Y.M., al-Zoghaibi, F., al-Haj, L., Dhalla, M., Zhou, A., Dong, B., Whitmore, M., Paranjape, J., Al-Ahdal, M.N., et al. (2003) RNase L mediates transient control of the interferon response through modulation of the double-stranded RNA-dependent protein kinase PKR. The Journal of biological chemistry, 278, 20124–20132.

56. Banerjee, S., Chakrabarti, A., Jha, B.K., Weiss, S.R. and Silverman, R.H. (2014) Cell-type-specific effects of RNase L on viral induction of beta interferon. mBio, 5, e00856–00814.

57. Oda, J.M., den Hartigh, A.B., Jackson, S.M., Tronco, A.R. and Fink, S.L. (2023) The unfolded protein response components IRE1alpha and XBP1 promote human coronavirus infection. mBio, 14, e0054023.

58. Nguyen, L.C., Renner, D.M., Silva, D., Yang, D., Parenti, N.A., Medina, K.M., Nicolaescu, V., Gula, H., Drayman, N., Valdespino, A. et al. (2022) SARS-CoV-2 Diverges from Other Betacoronaviruses in Only Partially Activating the IRE1alpha/XBP1 Endoplasmic Reticulum Stress Pathway in Human Lung-Derived Cells. mBio, 13, e0241522.

59. Thoresen, D.T., Galls, D., Gotte, B., Wang, W. and Pyle, A.M. (2023) A rapid RIG-I signaling relay mediates efficient antiviral response. Mol Cell, 83, 90–104 e104.

60. Kayesh, M.E.H., Kohara, M. and Tsukiyama-Kohara, K. (2025) Effects of oxidative stress on viral infections: an overview. Npj Viruses, 3, 27.

