## Supplementary Information for "RNase L Regulates Antiviral Responsiveness through Cleavage of XBP1 mRNA"

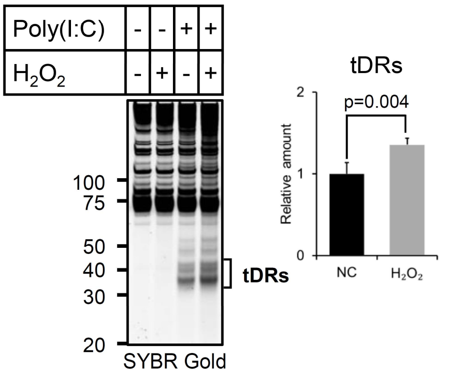


**Supplementary Figure S1. Oxidative inhibition of RTCB by $H_2O_2$ enhances Poly(I:C)-induced tDR production in A549 cells.**

tDR levels were evaluated by SYBR Gold staining 2 hours after $H_2O_2$ treatment, which was added 4 hours after Poly(I:C) transfection (as shown in Figure 1B). The amounts of tDRs calculated by densitometry are also shown (n=3).

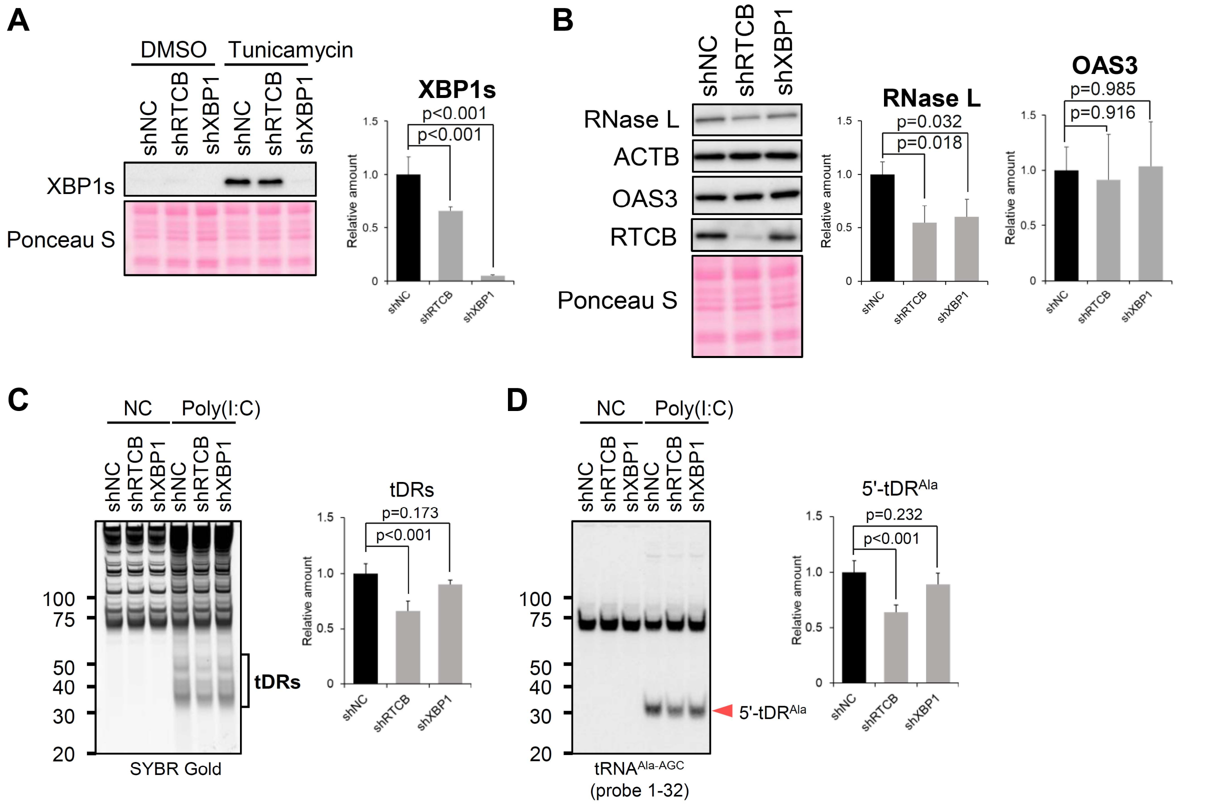


**Supplemental Figure S2. XBP1s acts as a positive regulator of the RNase L pathway in A549 cells.**

(A) RTCB/XBP1 knockdown significantly decreases tunicamycin-induced XBP1s production in A549 cells. Relative amounts of tunicamycin-induced XBP1s calculated by densitometry are also shown (n=4). (B) XBP1s positively regulates RNase L expression in A549 cells. Western blot analysis of RNase L, ACTB, OAS3 and RTCB. Relative amounts of RNase L (n=3) and OAS3 (n=4) calculated by densitometry are also shown. (C, D) RTCB/XBP1 knockdown decreases RNase L-induced tDRs in A549 cells. (C) SYBR Gold staining and (D) Northern blot analysis of $tRNA^{Ala\text{-}AGC}$. Relative amount of tDRs and $5'\text{-}tDR^{Ala}$ calculated by densitometry are also shown (n=4 each).

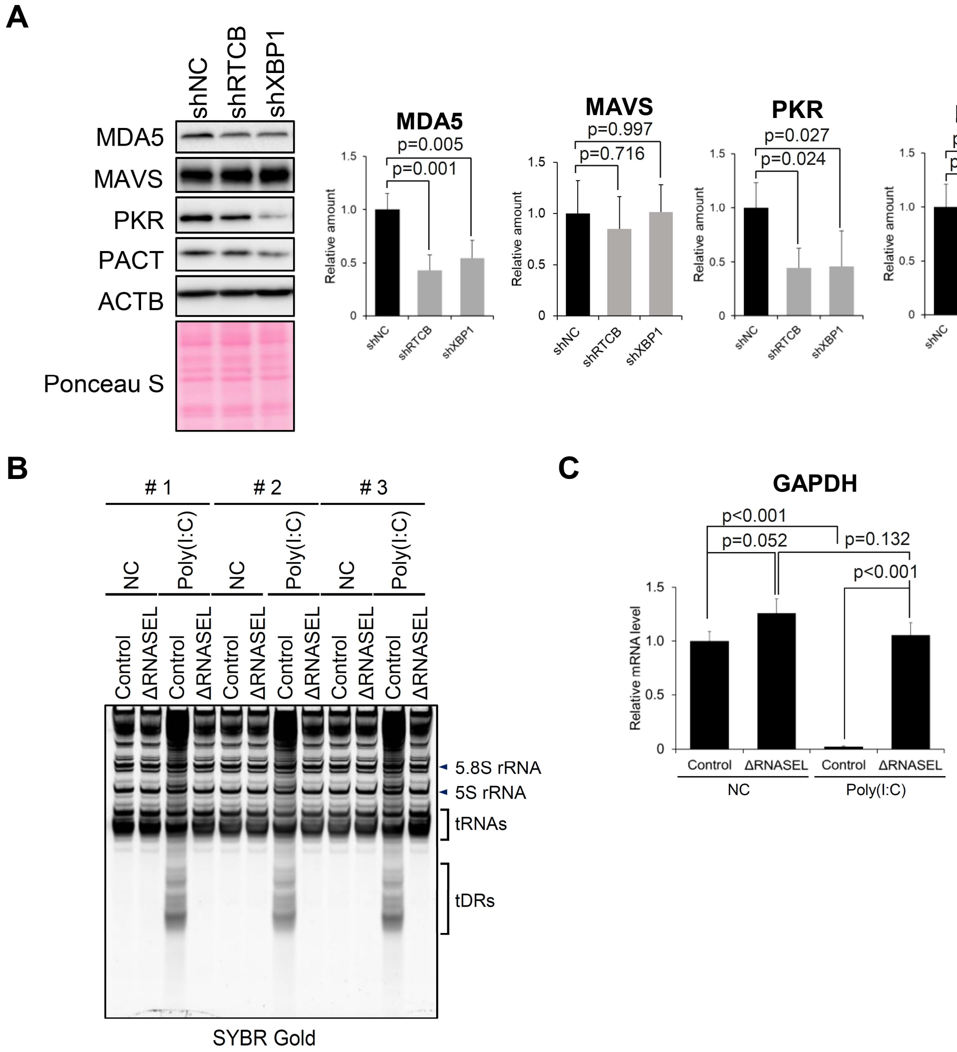


**Supplemental Figure S3. XBP1s positively regulates all three antiviral response pathways in A549 cells.**

(A) RTCB/XBP1 knockdown reduces the expression of MDA5, PKR and PACT in A549 cells. Western blotting for MDA5, MAVS, PKR, PACT and ACTB. Relative amounts of MDA5, MAVS, PKR and PACT calculated by densitometry are also shown (n=4 each). (B) RNase L-mediated RNA degradation 6 hours after transfection with 2 µg/mL Poly(I:C). (C) GAPDH mRNA levels under the same conditions, quantified based on equal total RNA input. Means and standard deviation were obtained from three independent experiments.

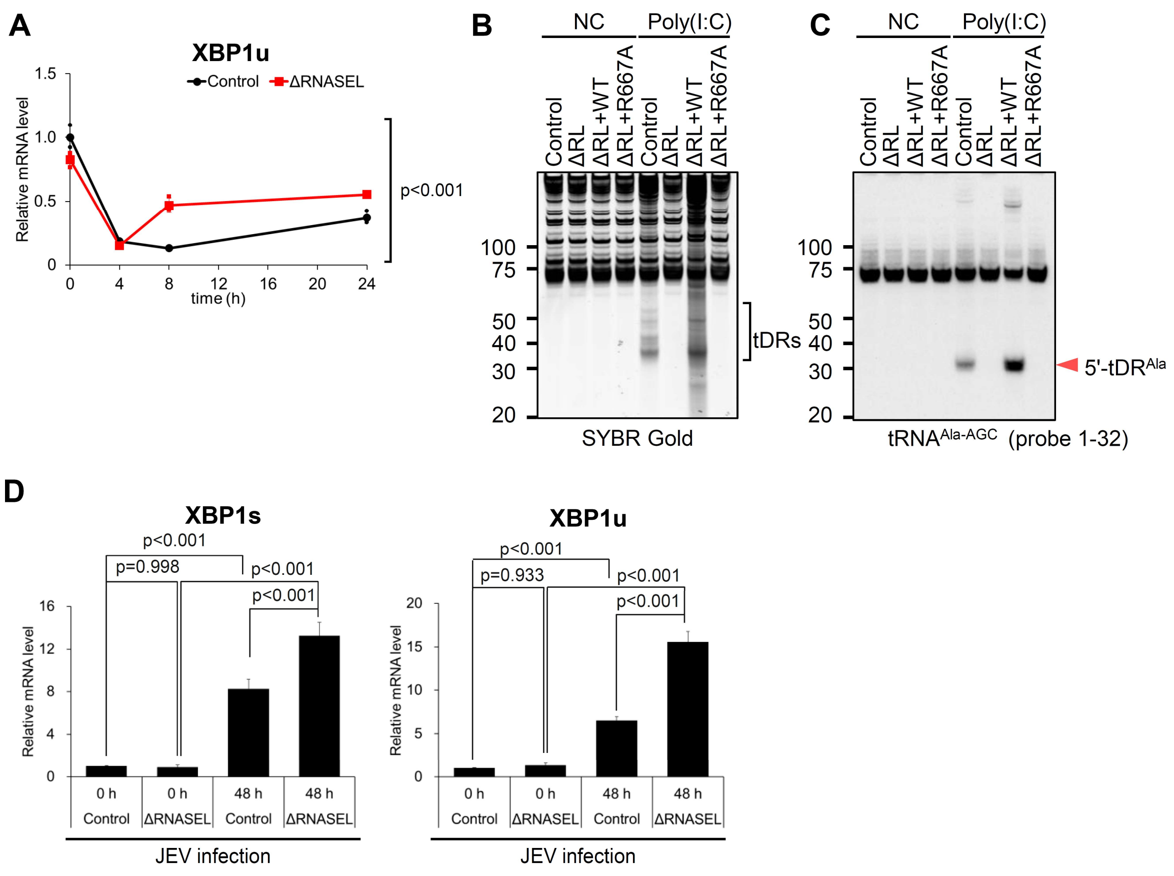


**Supplemental Figure S4. RNase L reduces XBP1 mRNA levels and suppresses XBP1s production (related to Figure 4).**

(A) Time-dependent induction of XBP1u mRNA in the Poly(I:C)-thapsigargin combination model (n=3 each). (B, C) Reconstitution of wild-type RNase L in ΔRNASEL cells restored Poly(I:C)-induced tDR production. (B) SYBR Gold staining and (C) Northern blot analysis of $tRNA^{Ala\text{-}AGC}$ at 6 hours after transfection. (D) Additional data related to Figure 4H. Relative XBP1u and XBP1s mRNA expression normalized to GAPDH mRNA by the ΔΔCT method.

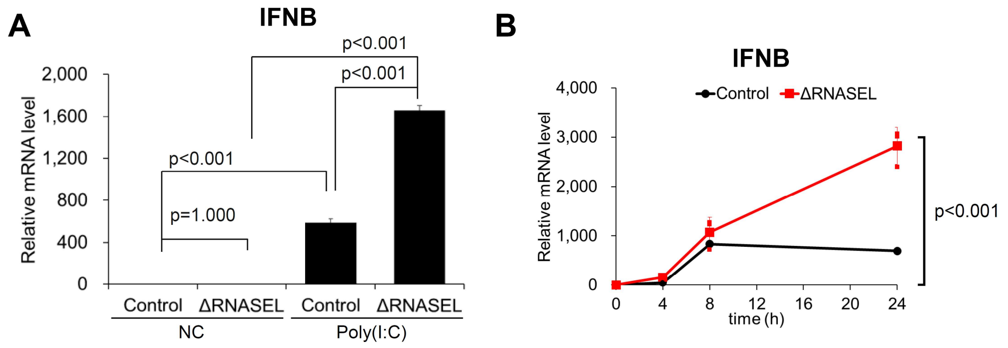


**Supplemental Figure S5. RNase L limits IFNB mRNA induction by reducing XBP1s production (related to Figure 5).**

(A) Relative IFNB mRNA expression levels normalized to GAPDH mRNA by the ΔΔCT method at 24 hours in the Poly(I:C)-thapsigargin combination model. (B) Time course of IFNB mRNA induction, normalized to GAPDH mRNA and calculated by the ΔΔCT method. Data are from three independent experiments.

| Antibodies | Source | Identifier |
|---|---|---|
| Rabbit anti-OAS3 | Proteintech | Cat #21915-1-AP |
| Mouse anti-RNase L | Santa Cruz Biotechnology | Cat #sc-74405 |
| Rabbit anti-XBP1s | abcam | Cat #ab220783 |
| Mouse anti-HSPC117/FAAP (RTCB) | Santa Cruz Biotechnology | Cat #sc-393966 |
| Rabbit anti-PKR [EPR19374] | abcam | Cat #ab184257 |
| Rabbit anti-PKR (phospho T446) [E120] | abcam | Cat #ab32036 |
| Rabbit anti-PACT (D9N6J) | Cell Signaling Technology | Cat #13490S |
| Rabbit anti-MDA-5 (D74E4) | Cell Signaling Technology | Cat #5321 |
| Mouse anti-MAVS (E-3) | Santa Cruz Biotechnology | Cat #sc-166583 |
| Mouse anti-DYKDDDDK tag | Proteintech | Cat #66008-4-IG |
| Mouse anti-Beta Actin | Proteintech | Cat #66009-1-IG |
| HRP-conjugated Goat anti-mouse IgG | Jackson Immunoresearch Laboratories | Cat #115-035-062 |
| HRP-conjugated Goat anti-rabbit IgG | Jackson Immunoresearch Laboratories | Cat #111-035-144 |
| Anti-rabbit IgG, HRP-linked Antibody | Cell Signaling Technology | Cat #7074S |

**Supplemental Table 1. List of antibodies used in this study.**

| shRNAs | | Sequences (5' to 3') |
|---|---|---|
| shNC | Top Strand | ccggGCATTCACTTGGATAGTAActcgagTTACTATCCAAGTGAATGCttttt |
| | Bottom Strand | aattaaaaaGCATTCACTTGGATAGTAActcgagTTACTATCCAAGTGAATGC |
| shRTCB | Top Strand | ccggGGAATTGTTCATCGATCTActcgagTAGATCGATGAACAATTCCttttt |
| | Bottom Strand | aattaaaaaGGAATTGTTCATCGATCTActcgagTAGATCGATGAACAATTCC |
| shXBP1 | Top Strand | ccggGAGAATTCCTCTATTTGTTCActcgagTGAACAAATAGAGGAATTCTCttttt |
| | Bottom Strand | aattaaaaaGAGAATTCCTCTATTTGTTCActcgagTGAACAAATAGAGGAATTCTC |

**Supplemental Table 2. Sequences of DNA oligos inserted to pLKO.1 vector for shRNA-mediated knockdown.**

| Primers | | Sequences (5' to 3') |
|---|---|---|
| FLAG-RNASEL | Forward | cgcGGATCCaccgtcatggattacaaggatgacgacgataaggagagcagggatcataac |
| | Reverse | CGTgaattctcagcacccagggctg |
| PAM mutagenesis | Forward | Agctgcagtggaagacaatcacttg |
| | Reverse | cttctaccgctggaggacgtgg |
| R667A mutagenesis | Forward | GCCaatttgggagaacacattga |
| | Reverse | gatgaactttagcagatcacc |
| XBP1s over expression | Forward | acgcggatccGCCACCatggtggtggtggcagccgc |
| | Reverse | gcgtCTCGAGttagacactaatcagctggggaaaga |

**Supplemental Table 3. Sequences of primers for PCR used in this study.**

| Probes | | |
|---|---|---|
| | Target Position | Sequences |
| tRNA$^{Ala-AGC}$ | 1-32 | 5'-AAGCACGCGCTCTACCACTGAGCTACACCCCC-3' |

**Supplemental Table 4. Sequences of DNA oligo probes for Northern blotting used in this study.**

| Primers | | Sequences (5' to 3') |
|---|---|---|
| OAS1 | Forward | CAAGCTCAAGAGCCTCATCC |
| | Reverse | TGGGCTGTGTTGAAATGTGT |
| OAS3 | Forward | TGAATTTCTCCAGCCCAACCG |
| | Reverse | TGGCTGAAGAGCCACCCTTG |
| RNase L | Forward | ACAAGTGGACGACTAAGATTAATGA |
| | Reverse | AGCAGATCACCCACAGTGTT |
| IFNB | Forward | AAACTCATGAGCAGTCTGCA |
| | Reverse | AGGAGATCTTCAGTTTCGGAGG |
| total XBP1 | Forward | GAAGGCGCTGAGGAGGAAA |
| | Reverse | CCAGCTCACTCATTCGAGCC |
| XBP1u | Forward | GGCGGAAGCCAAGGGGAAT |
| | Reverse | GCAGAGGTGCACGTAGTCTG |
| XBP1s | Forward | GCTGAGTCCGCAGCAGG |
| | Reverse | GTCCAGAATGCCCAACAGGA |
| GAPDH | Forward | GTCGGAGTCAACGGATTTGG |
| | Reverse | ATGGAATTTGCCATGGGTGGA |

**Supplemental Table 5. Sequences of primers for qPCR used in this study.**
